# Chitinase-3-like protein 1 decodes chitosan acetylation patterns into toll-like receptor 2 signaling through heparan sulfate

**DOI:** 10.64898/2026.06.15.732264

**Authors:** Alexandra Großdorf, Sina Nabil, Carlotta Imelmann, Margareta J. Hellmann, Jurij Froese, Ehab El-Awaad, Tuan Anh Tran, Stefan Cord-Landwehr, Nick Rähse, Önder Kurc, Fuzhu Yang, Hung-Yu Chen, Evelyne Gout, Gabriela Rosca, Marion Kusche-Gullberg, Martina Delbianco, Jonathan Cramer, Holger Gohlke, Romain R. Vivès, Alexander N.R. Weber, Hans Merzendorfer, Anayancy Osorio-Madrazo, Kay Grobe, Bruno M. Moerschbacher, Christian Gorzelanny

**Affiliations:** Department of Dermatology und Venereology, University Medical Center Hamburg-Eppendorf, Hamburg, Germany; Institute for Biology and Biotechnology of Plants, University of Münster, Münster, Germany; Institute of Physiological Chemistry and Pathobiochemistry, University of Münster, Münster, Germany; Department of Chemistry-Biology, University of Siegen, Siegen, Germany; Department of Pharmacology, Faculty of Medicine, Assiut University, Egypt; Laboratory of Organ Printing, University of Bayreuth, Bayreuth, Germany; Heinrich Heine University Düsseldorf, Faculty of Mathematics and Natural Sciences, Institute for Pharmaceutical and Medicinal Chemistry, Düsseldorf, Germany; Department of Biomolecular Systems, Max-Planck-Institute of Colloids and Interfaces, Potsdam, Germany; University Grenoble Alpes, CNRS, CEA, IBS, Grenoble, France; Department of Biomedicine and Centre for Cancer Biomarkers, University of Bergen, Bergen, Norway; Forschungszentrum Jülich, Institute of Bio-and Geosciences (IBG-4: Bioinformatics), Jülich, Germany; Department of Immunology, University of Tübingen, Tübingen, Germany

## Abstract

Chitinase-3-like protein 1 (CHI3L1), which is associated with a wide range of inflammatory diseases, lacks chitinase activity but retains the ability to bind chitin and chitosan. Chitin is a major component of fungal cell walls, whereas chitosan is used in biomedicine. In addition to chitosan, CHI3L1 has been proposed to interact with heparan sulfate (HS), a highly sulfated glycosaminoglycan on mammalian cell surfaces. Here, we investigated how interactions with chitosan and HS regulate the pro-inflammatory activity of CHI3L1. Mapping of the chitin-binding cleft revealed preferential binding of CHI3L1 to chitosans with a regular acetylation pattern that, together with CHI3L1, promoted toll-like receptor 2 signaling. We further identified a dominant HS-binding site that recognizes a distinct HS sulfation code containing a coherent motif of N- and 6-O-sulfations. Mutation of this HS-binding site or impaired HS biosynthesis prevented CHI3L1 accumulation at the cell surface and abolished CHI3L1-mediated cell activation. Together, our findings establish HS as a critical co-receptor for CHI3L1 and reveal a pro-inflammatory cross-talk between HS, CHI3L1, and chitosan that may contribute to host defense against fungal pathogens and responses to chitosan-based biomaterials. These findings identify HS- and chitosan-dependent CHI3L1 signaling as a potential target for modulating inflammatory responses.

## Introduction

Chitin is one of the most abundant polysaccharides on Earth and a crucial structural component of crustacean shells, insect exoskeletons, and fungal cell walls. It is a homopolymer composed of β-1,4-linked *N*-acetylglucosamine (GlcNAc). Partial deacetylation of chitin yields chitosan, a material of high bioeconomic interest. Chitosan is used in diverse applications such as wastewater treatment^1^, cosmetics^2^, food production^3^ and medicine.^4^ Common medical applications of chitosan include the generation of scaffolds for tissue engineering, drug delivery systems, and wound dressings. While chitin is generally considered pro-inflammatory, chitosan has long been regarded as immunologically inert. Previous studies have identified several putative chitin receptors that can trigger innate immune responses in mammals, including LysMD3, FIBCD1, and toll-like receptor 2 (TLR2).^5–7^ TLR2 is abundantly expressed in various cell types, including immune and epithelial cells. Upon ligand binding, TLR2 forms either homodimers or heterodimers together with TLR1 or TLR6. In addition to chitin, TLR2 recognizes a wide range of pathogen-associated molecular patterns (PAMPs), including zymosan and peptidoglycan.^8^ Activation of TLR2 signaling induces the expression of pro-inflammatory cytokines such as interleukin-8 through activation of the transcription factor NF-κB. As chitin is not produced by mammals, it is absent under physiological conditions and only encountered through external sources such as fungal pathogens or insects. Nevertheless, mammalian cells express chitinases capable of recognizing and processing chitin. Chitinases bind chitin within a defined chitin-binding cleft, where GlcNAc residues occupy specific subsites. Structural and biochemical analyses of CHIT1 have shown that some of these subsites can also accommodate glucosamine (GlcN), allowing binding and hydrolysis of chitosan.^9^ These findings challenge the previous assumption that chitosan is intrinsically immunologically inert. Indeed, more recent studies indicate that the immunological activity of chitosan depends on its degree of acetylation (DA) and, importantly, its pattern of acetylation (PA).^10,11^ We previously demonstrated that chitosans with a high DA promote macrophage activation more efficiently than those with a low DA.^9^ Furthermore, chitosans with a more regular PA induce stronger pro-inflammatory responses than those with a random PA.^10^

The mammalian chitinase family comprises six members, including the catalytically active enzymes CHIT1 and AMCase, as well as the enzymatically inactive chitinase-like proteins chitinase 3-like protein 1 (CHI3L1, also known as YKL-40), chitinase 3-like protein 2 (CHI3L2, also known as YKL-39), and oviductin-specific glycoprotein.^12^ Although the chitinase-like proteins have lost their enzymatic activity, they retain the ability to bind chitin. While CHIT1 has been mechanistically linked to chitosan-induced inflammation, the potential role of CHI3L1 in this context remains unclear.^9^ Independent of its chitin-binding capacity, CHI3L1 is present in blood plasma at low levels but its concentration increases under inflammatory conditions. Therefore, CHI3L1 has been proposed as a biomarker for several inflammatory diseases, including atopic dermatitis, arthritis, and inflammatory bowel disease.^13–15^ These conditions affect epithelial tissues of the skin and intestine, respectively, where CHI3L1 expression is markedly upregulated during inflammation, suggesting a potential pathophysiological role.

Earlier studies proposed that CHI3L1 acts as a ligand for IL-13Rα2, a receptor described as a decoy that binds IL-13 without triggering an immune response. However, binding of CHI3L1 to IL-13Rα2 can induce intracellular signaling through interaction with the transmembrane protein TMEM219. More recent studies also implicate CD44 in CHI3L1 signaling. Interestingly, although CD44 is primarily known as a hyaluronan receptor, its interaction with CHI3L1 is likely mediated by a heparan sulfate (HS) chain covalently attached to CD44.^16^ HS is a sulfated glycosaminoglycan abundantly present on the surface of mammalian cells. In addition to CD44, HS chains are commonly attached to the transmembrane proteoglycans syndecans (SDCs) or to glypicans. Previous studies have shown that binding of CHI3L1 to SDC1 promotes tumor angiogenesis. Although the chitin-binding activity of CHI3L1 has been studied previously, it remains unclear whether co-assembly of chitin or chitosan with HS modulates the pro-inflammatory activity of CHI3L1. HS frequently acts as a co-receptor for growth factors, cytokines or morphogens. Protein–HS interactions are strongly influenced by the degree and pattern of HS sulfation. Alterations of HS sulfation patterns are linked to the development of various diseases such as cancer^17,18^, Alzheimer’s disease^19^ and inflammatory bowel disease^20^. HS biosynthesis, including HS sulfation occurs in the Golgi apparatus and involves more than 20 enzymes.^21^ HS polymer elongation is catalyzed by a complex of exostosin-1 (EXT1) and exostosin-2 (EXT2), generating a chain of alternating GlcNAc and glucuronic acid (GlcA) residues. Subsequent *N*-deacetylation and *N*-sulfation of GlcNAc residues to form GlcNS are catalyzed by *N*-deacetylase/*N*-sulfotransferases (NDSTs). Additional sulfate groups are introduced by HS 2-O-, 3-O-, and 6-O-sulfotransferases (HS2ST1, HS3STs, and HS6STs). Further remodeling of HS fine structure and sulfation pattern is provided by Sulfs, a family of extracellular endosulfatases.^22^ Because sulfation is neither obligatory nor uniform across cell types, HS chains vary in length as well as in their degree and pattern of sulfation.

Here, we propose that CHI3L1 acts as a molecular shuttle that captures chitin or chitosan in the extracellular space and delivers it to cell surface-exposed TLR2. Cell surface HS may contribute to this process by promoting the accumulation of CHI3L1, thereby amplifying pro-inflammatory signaling through TLR2.

## Material and methods

### Fluorescence spectroscopy

Intrinsic protein fluorescence was measured using a multimode microplate reader (Spark® Tecan, Männedorf, Switzerland). CHI3L1 at a concentration of 1 µM was mixed with chitin hexamer (A6), chitosans or heparins at indicated concentrations in 25 mM Tris HCl and 150 mM NaCl buffer at pH 7.4 in black 96-well plates (Brand, Wertheim, Germany). The final volume was 100 μL per well. Intrinsic fluorescence was excited at 290 nm and an emission spectrum from 320 to 500 nm was recorded. For quantification, the fluorescence intensity at 340 nm was analyzed.

### Activity measurement of activated CHIT3L1

The chitinase substrate 4-methylumbelliferyl β-D-N,N’-diacetylchitobioside hydrate (Sigma, St. Louis, MO, USA) was diluted in 100 mL McIlvaine buffer (100 mM citric acid, 200 mM sodium phosphate, pH 5.2) at a final concentration of 20 µM. The enzyme reaction was started by adding 5 µL of HEK293F cell supernatant containing the activated CHI3L1. The reaction was stopped after 20 min at 37 °C by the addition of 200 µL of a 300 mM glycine-NaOH buffer, pH 10.5. Fluorescence was measured at 360 nm excitation and 450 nm emission in a fluorescence spectrometer (Infinite 200Pro, Tecan).

### TLR2 reporter assay

HEK-Blue™ hTLR2 stimulation was performed as previously described.^5^ Briefly, cells were cultured in T75 cell culture flasks in DMEM supplemented with 10% fetal calf serum, 100 mg/mL hygromycin and 50 mg/mL zeocin at 37 °C and 5% CO_2_ and passaged every three to four days. For stimulation, the cells were washed with PBS and then incubated with 5 mL PBS for 3 min at 37 °C. The cells were detached from the T75, counted, and centrifuged. The cells were then resuspended in HEK-Blue™ detection medium and seeded into a 96-well plate at a density of 50,000 cells per well. Cells were stimulated by chitosan in the absence or presence of CHI3L1 in the serum free medium, OptiPRO™ SFM. The synthetic TLR2 ligand PAM3CSK4 (1 μg/mL) was used as a positive control, and unstimulated cells served as a negative control. The plate was incubated for 24 h at 37 °C and 5% CO₂. Absorbance at 620 nm was measured using a microplate reader (Spark®, Tecan). Where indicated cells were pretreated with recombinant SULF2 for 1 h at 37 °C in HEK-Blue™ detection medium containing CaCl_2_ (1 mM).

### Endotoxin test

Chitosans and recombinant CHI3L1 were tested for endotoxin contamination using the Pierce™ Chromogenic Endotoxin Quant Kit (Thermo Fisher Scientific, USA) according to the manufacturer’s instructions.

### Enzymatic hydrolysis of polymers

Chitosan polymers (DA 15 & DA 45%) were hydrolyzed at a concentration of 1 g/L in ammonium acetate buffer (100 mM, pH 5) with rCHI3L1 (0.15 mg/L, respectively). Live digestions were performed as previously described^23^. Briefly, 80 μL of the rCHI3L1 polymer mixture were prepared in triplicates for each tested enzyme-substrate combination and incubated at 37 °C. After 180 min and 24 h of incubation, 30 μL of the samples was filtered through centrifugal filters (modified PES, 3 kDa cutoff; VWR, Germany) to separate oligomer products to be used for subsite preference analysis from enzyme and remaining polymer. Prior to SEC-RI-MS analysis (see SEC-RI-MS section), the enzyme reaction was stopped by addition of 1.1 μL HCl (250 mM).

### SEC-RI-MS

The analysis of hydrolysates by SEC-RI-MS was performed as described previously.^23^ Briefly, samples containing 3 μg of chitosan were separated with an ACQUITY UPLC Protein BEH SEC column with a pore size of 125 Å (Waters Corporation, USA), followed by RI detection (ERC RefractoMax 524; Thermo Fisher Scientific, USA) and electrospray ionization MS detection (amaZon speed; Bruker, Germany). MS data was analyzed with Data Analysis 4.1 (Bruker, Germany) and an in-house Python script based on the module pymzML^24^, RI data was evaluated using Data Analysis 4.1 (Bruker, Germany) and OriginPro 2023 (OriginLab, USA).

### Subsite preference analysis

Subsite preference analysis was performed as previously described.^25^ Briefly, the filtered digested polymers (see section enzymatic hydrolysis of polymers) were dried in a vacuum concentrator (30 °C) and subsequently dissolved in 50 μL 1:1 NaHCO_3_ (50 mM): MeOH. Two times, 1 μL of [^2^H_6_]-acetic anhydride (Sigma-Aldrich, Steinheim, Germany, USA) was added, each time followed by rapid mixing and incubation at 30 °C and 1200 rpm for 30 min. The *N*-acetylated oligomers were again dried in a vacuum concentrator (30 °C) and subsequently dissolved in different volumes of MilliQ water, based on the amount of small digestion products analyzed by SEC-RI-MS: 10 μL for rCHI3L1 on DA 15% and 48% chitosan for both time points. For MS1 analysis, 1.56 μL of these dissolved *N*-acetylated samples was mixed with 1.44 μL MilliQ containing 24 ng of each internal chitin standard of DP 1–6, which are double-isotopically labeled via *N*-acetylation with [^2^H_6_; ^13^C_4_]-acetic anhydride (Sigma-Aldrich, Steinheim, Germany, USA). A volume of 2 μL (including 16 ng of each standard) was analyzed by hydrophilic interaction liquid chromatography (HILIC) coupled to MS1. For MS2 analysis, different volumes of the dissolved *N*-acetylated samples corresponding to 10 μg chitosan based on the polymer substrates were dried in a vacuum concentrator (30 °C). The samples were then dissolved in 10 μL H_2_^18^O (Sigma-Aldrich, Steinheim, Germany, USA), incubated overnight at 70 °C, and 2 μL were analyzed by HILIC-MS2.

HILIC was performed with an ACQUITY Premier UPLC System (Waters, USA) using an Acquity UPLC BEH Amide column (1.7 μm, 2.1 mm × 50 mm; Waters, USA) and a VanGuard precolumn (1.7 μm, 2.1 mm × 5 mm; Waters, USA) with a column oven temperature of 40 °C. The flow rate was set to 0.4 mL/min and the separation was performed over 10.5 min with the following gradient elution profile: 100% (v/v) A (80:20 acetonitrile:H_2_O with 10 mM NH_4_HCO_2_ and 0.1% (v/v) HCOOH) for 0.0–1.0 min, linear gradient to 20% (v/v) B (20:80 acetonitrile:H_2_O with 10 mM NH_4_HCO_2_ and 0.1% (v/v) HCOOH) from 1.0 to 7.5 min, linear gradient to 75% (v/v) B from 7.5 to 8.5 min, 75% (v/v) B for 8.5 to 9.0 min, linear gradient to 100% (v/v) A from 9.0 to 9.2 min, 100% (v/v) A for 9.2–10.5 min. The detection of separated oligomers was performed with a Synapt XS HDMS 4k mass spectrometer (Waters, USA). MS1 measurements were conducted in positive resolution mode with normal dynamic range sensitivity, with a mass range of m/z 200 to 1500 and a scan time of 0.25 s. To improve mass accuracy, leucine encephalin was injected in 10 s-intervals for 1 s for lock spray calibration. The source capillary voltage, sampling cone, and source offset were set to 3 kV, 35, and 20, respectively. The source and desolvation temperatures were 80 °C and 250 °C, respectively. Flow rates of 0 L/h cone gas and 500 L/h desolvation gas were used, and the nebulizer pressure was set to 5.8 bar. MS2 measurements were conducted in TOF MS/MS mode and each oligomer containing at least one natural and one isotopically labeled acetyl group was individually isolated using the quadrupole of the MS system with a LM resolution of 15 to enable the isolation and fragmentation of mono-isotopic peaks. The collision energy in the trap was set to 18 V for dimers, 20 V for trimers, 22 V for tetramers, 24 V for pentamers, and 26 V for hexamers. MS data was analyzed using MassLynx V4.2 (Waters, USA) and an in-house Python script based on the module pymzML^24^.

### Preparation of starting structures

The chitohexaose-(AAAAAA)-bound CHI3L1 crystal structure (PDB ID: 1HJW)^26^ was prepared using the Protein Preparation Workflow^27^ (default settings) in Maestro (Schrödinger Release 2024-4: Maestro, Schrödinger, LLC, New York, NY, 2024). N- and C-terminal residues were capped with ACE and NMA residues, respectively. The pK_a_ values and protonation states were predicted for pH 7.4, with all titratable residues subsequently assigned to their dominant protonation state. The N-linked glycan, at N60, visible in the crystal structure, was removed during system preparation. Starting structures for the respective CHI3L1-COS complexes (AAAAAD, AAAADA, AAADAA, AADAAA, ADAAAA, DAAAAA) monodeacetylated at the GlcNAc moiety were built using Maestro, by removing the acetyl group attached to the amino functionality at the C2 position to obtain the GlcN unit. Because the amino groups of the GlcN units are mainly unprotonated at pH 7.4, these ligands remained uncharged.^28^ The automated tool PrepareForLeap^29^ was used to map the carbohydrate ligands for use with the GLYCAM force field. The systems were solvated in a cubic box of OPC water^30^ such that the distance between the boundary of the box and the closest solute atom was at least 12 Å. To obtain a neutral system, we complemented the solution with explicit counterions (0.15 M NaCl) that replaced solvent molecules, according to the SPLIT method.^31^ Topology files were built using the tLEaP module in AmberTools24.^32^ The ff19SB force field for proteins^33^ was used in combination with the GLYCAM06 force field for the carbohydrates^34^ and the Li and Merz 12-6 ion parameters for Na+ and Cl– ions.^35,36^ For non-modified, terminal and inner β-D-glucosamine units, the corresponding prep file, provided by David Thieker in the GLYCAM mailing list (Mar 6, 2017), was used.

### Molecular dynamics simulations

All molecular dynamics (MD) simulations were performed with the AMBER24 suite of molecular simulation programs^32^ using the mixed-precision (SPFP) GPU-accelerated implementation of PMEMD.^37,38^ An adjusted protocol for the setup of simulations was used as described previously.^39^ All subsequent steps were repeated 12 times (different random seed for every run) for each complex to yield 12 independent simulations per system. Initially, the energy of the system was minimized by relieving unfavorable contacts of the water molecules. Solvent molecules were first minimized for 2,500 steps using the steepest descent algorithm, followed by 2,500 steps of minimization with the conjugate gradient algorithm. Remaining structural elements involving the protein and the ligand were restrained by applying harmonic positional potentials with force constants of 2.5-10.0 kcal mol^−1^ Å^−2^ (**Table 1**). Afterward, the system was heated over 50 ps from 100 K to 300 K in the NVT ensemble. Finally, the system was simulated in the NPT ensemble to adjust the density to 1 g cm^-3^ and gradually reduce positional restraints in a stepwise manner (seven steps for a total simulation time of 950 ps). Pressure was maintained at 1 bar using isotropic position scaling with the Berendsen barostat^40^ with a pressure relaxation time of 1 ps. Temperature control at a target temperature of T = 300 K was maintained using Langevin dynamics^41^ with a collision frequency of γ = 1.0 ps^-1^. The SHAKE^42^ algorithm was applied to constrain covalent bonds involving hydrogen atoms. Periodic boundary conditions were applied in all directions. Long-range electrostatic interactions were computed using the Particle Mesh Ewald (PME) method,^43^ while a non-bonded cutoff of 9 Å was employed for both short-range electrostatics and van der Waals interactions. Hydrogen mass repartitioning was applied to allow the use of a 4 fs integration time step.^44^ The production phase was performed with the resulting system and consisted of 1 µs length, giving 12 µs of cumulative sampling data for all production runs per complex (84 µs in total). Coordinates were saved in a trajectory file at 100 ps intervals. Post-processing and analysis of the MD trajectories were carried out with CPPTRAJ^45^ of AmberTools24.^32^ Trajectories were visually inspected with VMD.^46^

**Table 1.** Simulation protocol. Characteristics of the minimization, thermalization, density adaptation, and production phases for the CHI3L1-COS systems.

| Process | No. of Steps | Algorithm | Restrained Selection and Force Constant<br>[kcal mol<sup>-1</sup> Å<sup>-2</sup>] |  |  |
| --- | --- | --- | --- | --- | --- |
|  |  |  | PBB<sup>a</sup> | PSC<sup>b</sup> | LIG<sup>c</sup> |
| Minimization | 2,500/2,500 | Steepest descent/Conjugate gradient | 10.0 | 5.0 | 2.5 |
| Process | Time [ps] | Ensemble/Time Step [fs] | Restrained Selection and Force Constant<br>[kcal mol<sup>-1</sup> Å<sup>-2</sup>] |  |  |
|  |  |  | PBB<sup>a</sup> | PSC<sup>b</sup> | LIG<sup>c</sup> |
| Thermalization | 50.0 | NVT/1.0 | 10.0 | 5.0 | 2.5 |
| Density adaptation 1 | 100.0 | NPT/2.0 | 5.0 | 2.5 | 2.5 |
| Density adaptation 2 | 100.0 | NPT/2.0 | 2.5 | 1.0 | 1.0 |
| Density adaptation 3 | 100.0 | NPT/4.0 | 1.0 | 0.5 | 0.5 |
| Density adaptation 4 | 100.0 | NPT/4.0 | 0.5 | 0.1 | 0.1 |
| Density adaptation 5 | 100.0 | NPT/4.0 | 0.1 | - | - |
| Density adaptation 6 | 150.0 | NPT/4.0 | 0.05 | - | - |
| Density adaptation 7 | 300.0 | NPT/4.0 | - | - | - |
| Production | 1.0 10<sup>6</sup> | NPT/4.0 | - | - | - |

<sup>a</sup> protein backbone atoms; <sup>b</sup> protein side-chain heavy atoms; <sup>c</sup> ligand atoms

### Computation of effective binding energies

Effective binding energies (ΔG_eff_), comprising the sum of gas-phase energies and solvation free energies,^47^ of fully and partially acetylated COS binding to CHI3L1 were calculated using the single-trajectory MM-PBSA approach^48^ as implemented in the MMPBSA.py module^49^ in AmberTools24^32^. The ante-MMPBSA.py module was used to create the compatible complex, receptor, and ligand topology files from the solvated topology file, employing mbondi2 radii for the solute atoms while removing counterions and water molecules. Conformational ensembles for all calculations were generated by extracting 1,000 snapshots per MD trajectory at regular intervals of 1 ns. The polar contribution to the solvation free energy was calculated using an ionic strength of 0.15 M, an internal dielectric constant (ε_int_) of 4 for the solute, and an external dielectric constant (ε_ext_) of 80 for the solvent. A solute dielectric constant of 4 has been used before to account for highly charged^50,51^ and highly solvent-accessible^52^ binding sites, as given in the case of CHI3L1. Configurational entropy contributions were omitted to reduce additional uncertainty in the computational analysis.^47,53^ The effective binding energies were averaged over the respective ensembles. The standard error of the mean (SEM) for a given system was calculated from the standard deviation (SD) of the 12 mean ΔG_eff_ values according to Equation 1:

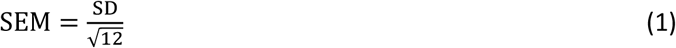

Relative effective binding energies (ΔΔG_eff_) were calculated by subtracting the effective binding energy of the fully acetylated ligand (ΔG_eff,FA_) from the effective binding energy of the partially acetylated ligand (ΔG_eff,PA_) (Equation 2).

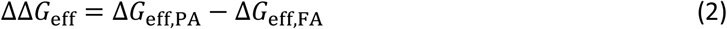

The total SEM of the relative effective binding energy (Equation 2) was calculated via Equation 3, following the law of error propagation:

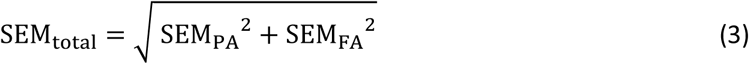

A two-sided Welch’s t-test was performed to probe if the average ΔG_eff_ values of the modified ligands (ΔG_eff,PA_) differ significantly from the native ligand (ΔG_eff,FA_), using SciPy.^54^ A p-value < 0.05 was considered significant.

### Microscale Thermophoresis (MST)

Protein-COS interactions were quantified using microscale thermophoresis (MST). The well-defined COS were prepared with Automated Glycan Assembly (AGA) from monosaccharide building blocks (BBs) following a previously reported procedure.^55^ All details concerning BBs synthesis, AGA modules, post-AGA manipulations, and characterization can be found in the Supplementary Information. MST experiments were performed on a NanoTemper Monolith NT.115 (Nanotemper, München, Germany). Cyanine 5-maleimide (Lumiprobe, Hannover, Germany) was employed for labeling the free Cys41 residue of CHI3L1 by incubating the protein with a 10-fold molar excess of the reactive dye in phosphate-buffered saline (10 mM Na_2_HPO_4_, 1.8 mM KH_2_PO_4_, 137 mM NaCl, 2.7 mM KCl) at pH 7.4 according to the manufacturer’s recommendations. Excess dye was removed by size-exclusion chromatography. The degree of labeling was calculated as 0.69. COS titration series were prepared in phosphate-buffered saline (10 mM Na_2_HPO_4_, 1.8 mM KH_2_PO, 137 mM NaCl, 2.7 mM KCl, 0.05% Tween-20) at pH 7.4 by serial dilution. After addition of the labeled protein to a final concentration of 20 nM, all samples were incubated for 15-25 min at room temperature to allow equilibration before loading into the capillaries. All data processing and curve fitting were performed using the manufacturer-supplied software and illustrated with Graph Prism 8 (GraphPad Software LLC, Boston, USA).

### Heparin-CHI3L1 binding assay

Heparin-coated plates (Bioworld, Dublin, USA) were used to measure the binding of CHI3L1 to heparin. CHI3L1/mutants (0.25 μM) were incubated with increasing amounts of soluble CHI3L1 ligands (chitin A6; unfractionated HS (UFH) (Biochrom AG, Berlin, Germany). Protein-ligand combinations were incubated in a 96-well plate for 1 h at room temperature in PBS. Plates were washed with 3x 300 µL 0.05% Tween-PBS (Fisher Scientific GmbH, Schwerte, DE) followed by blocking with 100 μL 1% bovine serum albumin (BSA, Sigma Aldrich, St. Louis, USA) for 30 min at room temperature, and the excess BSA was washed off with 3x 300 μL 0.05% Tween PBS. The primary antibody (100 μL, mouse anti-His Tag antibody (Origene, Rockville, USA) was diluted 1:1000 in PBS and incubated for 2 h at room temperature. After washing again with 3x 300 μL Tween PBS, detection was performed with the HRP-conjugated goat anti-mouse secondary antibody (Agilent, Santa Clara, USA) (100 μL, 1:5000 in PBS) for 20 min at room temperature.

After washing, 100 μL of HRP substrate (BD Biosciences, Franklin Lakes, USA) was incubated for 20 min and stopped with 50 μL H_2_SO_4_. Absorption was measured at 450 nm and 570 nm in the Spark® multimode microplate reader (Tecan, Männedorf, Switzerland), with 570 nm values subtracted from the 450 nm values as correction values. CHI3L1/mutants (0–1 μM) in PBS were applied as the standard curve.

### Chitosan dot blot and HS microarray assay

The chitosan dot blot assay was performed as previously reported with some modifications.^56^ Different chitosans (DA 0–50%) 1.5 μL were pipetted onto a nitrocellulose membrane in a dilution series (0–1000 ng/mL) (GE Healthcare, Munich, Germany). Membranes were dried at 70 °C for 30 min and subsequently blocked for 1 h with 5% (wt/v) BSA in Tris-buffered saline (TBS). After washing with TBS, membranes were incubated with CHI3L1 (1 µg/mL) for 1 h at room temperature. Excess protein was removed by washing 2x with TBS supplemented with 0.05% (v/v) Tween 20 and Triton X-100. The membrane was incubated with primary antibody directed against CHI3L1 (YKL-40) (AF2599; R&D Systems, Minneapolis, USA) for 2 h at room temperature, followed by washing steps. Detection was performed using a HRP-conjugated donkey anti-goat IgG-HRP secondary antibody (Dako, Glostrup, Denmark) for 2 h at room temperature. After washing, SuperSignal™ West Pico PLUS Chemiluminescent Substrate (Thermo Fisher Scientific, USA) was added, and signals were imaged by a chemiluminescence reader (Azure Biosystems, Dublin, USA).

The glycan microarray analysis was performed by Glycan Therapeutics (Raleigh, USA).

### Single-Molecule Force Spectroscopy (SMFS)

SMFS was performed as previously reported with some modifications.^57,58^ Briefly, we functionalized the AFM tip (BL-RC150VB-C1, Olympus, Tokio, Japan) covalently with a polyethylene glycol (PEG) linker (5000 Da, Nanocs, New York, USA) and CHI3L1/mutants (0.1 – 0.4 mg/mL). In parallel, we functionalized glass cover slides with a PEG linker, Streptavidin and Biotin-Heparin (1 mg/mL, Sigma-Aldrich, Steinheim, Germany). Prior to SMFS measurements (Nanowizard, JPK instruments, Berlin, Germany), spring constants (∼ 4 mN/m) of the cantilevers were calibrated using the thermal noise approach. Repeated force-distance cycles were performed at several loading rates in PBS at room temperature. After the tip contacted the cover glass, retraction was delayed for 1 s. Antibody detachment events were analyzed as previously reported (Equation 4):^59^

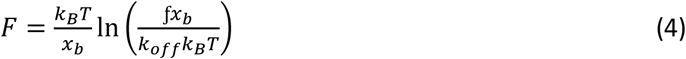

where *F* is the measured force required to detach the CHI3L1/mutant from the ligand, *k_off_* the off rate, *k_B_* the Boltzmann constant, T the temperature (293 K) and *x_b_* the depth of the energy barrier. Data analysis was performed with the JPK data processing software (version spm-5.0.131, JPK instruments) and an in-house Python script (version 3.9.7).

### QCM-D measurements

QCM-D was performed as previously reported.^60^ After functionalization of the sensor surface with lipid bilayers and heparin, QCM-D was measured with a QSense Analyser (Biolin Scientific, Gothenburg, Sweden) and SiO_2_-coated sensors (QSX303, Biolin Scientific). Measurements were performed at a working temperature of 23 °C with four parallel flow chambers and a peristaltic pump (Ismatec, Grevenbroich, Germany) with a flow rate of 20 µL per min. The normalized frequency shifts ΔF and the dissipation shifts ΔD, were measured at six overtones (i = 3, 5, 7, 9, 11, 13). Data were acquired with QSoft401 (version 2.7.3.883), and analyzed in OriginPro (version 10.2.0.188).

### GCI measurements

GCI measurements were carried out at room temperature using a Creoptix WAVEsystem (Malvern Panalytical, Switzerland) with PCH-NTA sensor chips. Prior to ligand immobilization, chip surface was conditioned by sequential injections of borate buffer (0.1 M sodium borate, 1 M NaCl, pH 9.0) and 0.25 M EDTA for 180 s each at a flow rate of 10 µL per min. For ligand immobilization, chip surface was activated with 0.5 mM NiSO_4_.6H_2_O for 120 s at 10 µL per min. CHI3L1 was injected onto one channel until the desired ligand density was reached, while the mutant variant CBmut CHI3L1 was immobilized on a reference channel of the same chip at a similar ligand density.

To analyze the interaction of CHI3L1 with COS analytes (DP5–10), dilution series of each analyte were prepared in phosphate-buffered saline (PBS, pH 7.4). Samples were injected from low to high concentrations at a flow rate of 30 µL per min for 60 s, followed by a 300 s dissociation phase under running buffer flow. Blank injections were included after every three sample injections for double referencing. Between measurements of different analytes, sensor surface was regenerated by injection of 0.5 M EDTA for 180 s at 10 µL per min. After regeneration, ligand immobilization was repeated as described above.

Sensorgrams were processed using Creoptix WAVEcontrol software (version 4.7.4). Apparent dissociation constants (K_D_) were determined after double referencing by fitting the data using conformational change binding model implemented in the software.

### HS characterization

HS chain length was analyzed as described previously.^61^ Briefly, subconfluent cells were metabolically labeled with 200 µCi/mL Na₂³⁵SO₄ (PerkinElmer) for 24 h. After incubation, the culture medium was collected and stored at −20 °C. Free glycosaminoglycan (GAG) chains were isolated from the trypsin fraction representing cell-surface and matrix-associated proteoglycans. Galactosaminoglycans were removed by digestion with chondroitinase ABC (Seikagaku) in Tris/HCl buffer (50 mM, pH 8.0) containing 30 mM sodium acetate and 0.1 mg/mL BSA. HS chain length was determined by gel chromatography on a Superose 6 HR 10/30 column (Amersham Biosciences) eluted with 0.5 M NH₄HCO₃, using previously established calibration standards.

### CHI3L1 binding to HS on genetically engineered cells

Binding of CHI3L1 to the mouse melanoma cell line B16F10 (ATCC #CRL-6475) was measured by flow cytometry. Cells were genetically engineered as previously described.^57^ Briefly, cells were cultured in DMEM (Sigma-Aldrich, Steinheim, Germany, St. Louis, USA) supplemented with 10% fetal bovine serum, 1% L-glutamine, and 1% penicillin/streptomycin. Cells were maintained at 37 °C in a humidified atmosphere containing 5% CO₂ and passaged upon reaching approximately 90% confluency. To generate pLenti-CMV_Blast_mKate2, the mKate2 sequence was amplified by PCR from the plasmid pLSS-mOrange2-mKate2 (Addgene #99868). The PCR product was digested with SalI and BamHI and subsequently ligated into the pLenti-CMV_Blast backbone (Addgene #17486) that had been digested with the same enzymes. For construction of pLenti-CMV_Blast_mKate2_gRNA, the plasmid pLenti-CMV_Blast_mKate2 was digested with BamHI and XhoI, and the resulting fragment was ligated with a gRNA expression cassette containing the hU6 promoter and gRNA scaffold sequence. Lentiviral particles were produced by co-transfecting HEK293 cells with pMDLg/pRRE (Addgene #12251), pRSV-Rev (Addgene #12253), pCMV-VSV-G (Addgene #8454), and the gRNA-containing construct pLenti-CMV_Blast_mKate2_gRNA using Lipofectamine 3000 (Thermo Fisher Scientific, Massachusetts, USA). B16F10 cells were transduced with the resulting lentivirus in the presence of Polybrene (Sigma-Aldrich, Steinheim, Germany) at a multiplicity of infection (MOI) of 1.0. Forty-eight hours after transduction, cells were selected with 7.5 µg/mL blasticidin for two weeks. For Cas9 expression, cells were transiently transfected with pIREShygroTO-Cas9 (BB)-2A-Puro using Lipofectamine 3000. Twenty-four hours post-transfection, puromycin selection (5 µg/mL) was applied for 48 h, after which puromycin was removed from the culture medium. Single-cell clones were isolated and verified by Sanger sequencing of the target region. Experiments were performed with pools of at least three independent clones.

For flow cytometry, cells washed and resuspended in PBS were incubated with CHI3L1/mutants (10 µg/mL) for 1 h on ice. As negative controls, the cells were incubated with 0.5% BSA instead of protein. After washing, the primary antibody (5 µg/mL) anti-CHI3L1 IgG (R&D Systems, Minneapolis, USA) in 0.5% BSA was added for 1 h. Alexa Fluor 647 anti-goat IgG (Invitrogen, Waltham, USA) (1:1000 in 0.5% BSA) was used as the secondary antibody for 30 min. To detect HS on the surface of the cells used, the cells were incubated with mouse anti-heparan sulfate antibody (Amsbio, Abingdon, UK) (1:4000) and Alexa Fluor 488 anti-mouse IgM (Jackson Immuno Research Laboratories Inc., West Grove, USA) (1:4000). Prior to the measurement, cells were washed and resuspended in 100 μL PBS. The measurement was performed on the NovoCyte Quanteon flow cytometer (Agilent, Santa Clara, USA). A maximum of 10,000 cells in 50 μL were measured within 1 min. The analysis was performed with the web tool Floreada.io.

### Preparation of defined chitin oligosaccharides

Fully acetylated chitin oligosaccharides were prepared by isolating soluble chitosan oligomers of single DPs and converting them to chitin by chemical *N*-acetylation. Briefly, a hydrolysate of low DA chitosan with a molecular weight below 3000 was fractionated by preparative size exclusion chromatography as described previously.^62^ Fractions of single DPs were pooled and freeze-dried. The integrity of the obtained chitosan oligomers was confirmed by HILIC-MS before chemical *N*-acetylation with acetic anhydride as described previously.^25^ The residual acetate was removed by repeated freeze-drying.

### Static light scattering

Aggregation of chitin oligosaccharides with sizes ranging from DP2 to DP10 was measured by static light scattering using the Spark® multimode microplate reader (Tecan, Männedorf, Switzerland). For this purpose, 2 µL of the chitin oligosaccharides (1 mg/mL) was pipetted onto a NanoQuant plate and light transmission was measured at a wavelength of 600 nm.

### Immunofluorescence staining

HEK-Blue™ hTLR2 reporter cells were seeded on an L-lysine-coated microscope slide (200,000 cells/slide) and cultivated overnight in DMEM supplemented with 10% fetal calf serum at 37 °C in 5% CO_2_. The next day, cells were washed with PBS and blocked with 1% BSA for 1 hour. Staining was performed in PBS containing 0.1% BSA with following antibodies: CHI3L1 (anti-CHI3L1 IgG primary antibody (R&D Systems, Minneapolis, USA) (1:13 in 0.5% BSA); Alexa Fluor 647 anti-goat IgG (Invitrogen, Waltham, USA) (1:1000 in 0.5% BSA)), heparan sulfate (anti-heparan sulfate antibody (Amsbio, Abingdon, UK) (1:4000) and Alexa Fluor 488 anti-mouse IgM (Jackson Immuno Research Laboratories Inc., West Grove, USA) (1:4000) and 4′,6-Diamidin-2-phenylindol-dihydrochlorid (Sigma-Aldrich, Steinheim, Germany)).

Slides were mounted with Fluoromount (Sigma-Aldrich, Steinheim, Germany) and dried overnight. Fluorescence images were generated using a fluorescence microscope (Observer z.1, Zeiss, Oberkochen, Germany) equipped with an Apotome module. Images were analyzed with the Zen software (version 3.11).

### Recombinant production of CHI3L1

For recombinant production of CHI3L1, the CHI3L1 gene was amplified from cDNA of human macrophages and cloned into the pIRES neo2 vector (Takara Bio, San Jose, USA). Point mutations were used to prevent chitin (W31A; W99A; W352A) or HS (R144A; R145A; K147A) binding or to activate (A138D; L140E) the protein. Mutagenesis for the chitin-binding mutant (CBmut CHI3L1) (mix 1) and the activated CHI3L1 (mix 2) was performed using the QuickChange Multi Site-Directed Mutagenesis Kit (Agilent, Santa Clara, USA), while Q5R Site-Directed Mutagenesis Kit (NewEngland Biolabs, Ipswich, USA) was used to produce the heparin-binding mutant (HepBmut CHI3L1) (mix 3). Mutagenesis mixtures 1 and 2 were digested with the enzyme DpnI, while mutagenesis mixture 3 was digested and ligated in one step according to the kit instructions. All mutagenesis mixtures were transformed into *E. coli* Mach T1 cells. Insertion of mutations was validated by Sanger sequencing. Plasmids were transfected into HEK293F cells and protein expressing cells were selected using geneticin.

HEK293F cells were maintained in DMEM (1% penicillin streptomycin, 1% L-glycine, 10% FCS) medium; for recombinant protein production, culture medium was replaced with serum-reduced Opti-MEM medium (Gibco) and cells were incubated at 37 °C with 5% CO_2_ for 24 h. Cell supernatant were centrifuged for 10 min at 4000 g and recombinant proteins equipped with a C-terminal His₆ tag were purified using a His-tag affinity column (5 mL HiTrap FF (Cytiva)) by liquid chromatography (ÄKTA go system, Cytiva) at room temperature, a constant flow rate of 1 mL/min and a linear imidazole gradient (20 CV, 0–500 mM). Protein-containing fractions were desalted using Zeba Spin desalting columns 7K MWCO 5 mL (Thermo Fisher) and buffered in PBS according to the manufacturer’s instructions. Protein purity was checked by polyacrylamide gel electrophoresis (NuPAGE™ 4 to 12%, Bis-Tris, 1.0–1.5 mm) mini protein gels. Gels were stained with Coomassie Brilliant Blue (CBB, Wako) and concentration of the purified proteins was determined using a BSA standard.

## Results

### CHI3L1 has distinct subsite preferences for GlcNAc and GlcN

Although active chitinases of glycosylhydrolase family 18 (GH 18), such as the CHI3L1-related CHIT1, preferentially degrade chitin, they were also able to cleave chitosans.^9^ Here, we tested the ability of CHI3L1 (**Figure 1A**, RCSB Protein Data Bank: 1NWR^63^) to bind to different chitosans with DAs ranging from 0% to 50% and a DP around 1000 (**Supplementary Figure 3C**). Binding affinity was highest for chitosan with a DA of 50% and lowest for DA 0% (**Figure 1B**). Although the binding to low-acetylated chitosans was reduced, our data suggest that some subsites within the chitin binding cleft tolerate GlcN. Previous analyses of active chitinases have shown that individual subsites within the chitin-binding cleft differ in their specificity for GlcNAc and GlcN: while some subsites display a strong preference for GlcNAc, others are more permissive and tolerate GlcN.^64^ Together, these subsite preferences determine which DA and PA are accepted or even preferred by a given enzyme. To define the subsite preferences of CHI3L1, we engineered an enzymatically active chitinase variant (CHI3L1 A138D/L140E) by introducing the mutations A138D and L140E via site-directed mutagenesis (**Figure 1C, D**). Activated CHI3L1 was able to cleave the fluorescent chitinase substrate 4-methyumbelliferyl chitobioside (**Supplementary Figure 1A**). In further experiments, we digested chitosans with a DA of 15% and 48% for 0.5 and 18 h and used size-exclusion chromatography to analyze the size of the generated oligosaccharides (**Figure 1E**). As expected for family 18 glycosylhydrolases, digestion of the DA 48% chitosan was more efficient than that of the DA 15% chitosan as indicated by the time-dependent decrease of the polymer peak and the simultaneous increase of oligomer abundance. The DA 15% chitosan remained largely unaffected. Oligomers produced from the DA 48% chitosan were further analyzed by mass spectrometry (**Figure 1F, Supplementary Figure 1B**). At the early time point of digestion (0.5 h), oligomers with a large fraction of GlcNAc (A units) and low fraction of GlcN (D units) such as A4D3, dominated. At the later time point, those oligomers disappear, while oligomers composed of A2D1 and A2D2 units appear. Although this finding already suggests that GlcN is accepted at certain subsites by the active CHI3L1, we further sequenced the produced oligomers by mass spectrometry to unravel distinct subsite preferences. We followed a previously established protocol and compared the subsite preference of CHI3L1 and CHIT1^64^, which shares high sequence similarity and identical amino acids in the distinct subsites, except for subsite –3 (**Figure 1G**). In line with the high structural similarity and as summarized in **Figure 1H**, the subsite preferences of CHI3L1 A138D/L140E closely resemble those reported for CHIT1.^64^ At subsite −1, GlcNAc was strongly preferred, whereas subsite +1 showed lower specificity and tolerated GlcN with comparable probability. Subsites −3, −2, +2 and +3 also preferred GlcNAc but readily accepted GlcN. The most pronounced difference between CHI3L1 and CHIT1 were found for subsite −3, which also showed the strongest deviation on the amino acid sequence level. CHI3L1 may contain additionally a −4 and +4 subsite, which we were not able to characterize due to technical limitations. However and as further outlined in the following section, we were able to map the −4 subsite by MD simulations.^64^

**Fig. 1:**
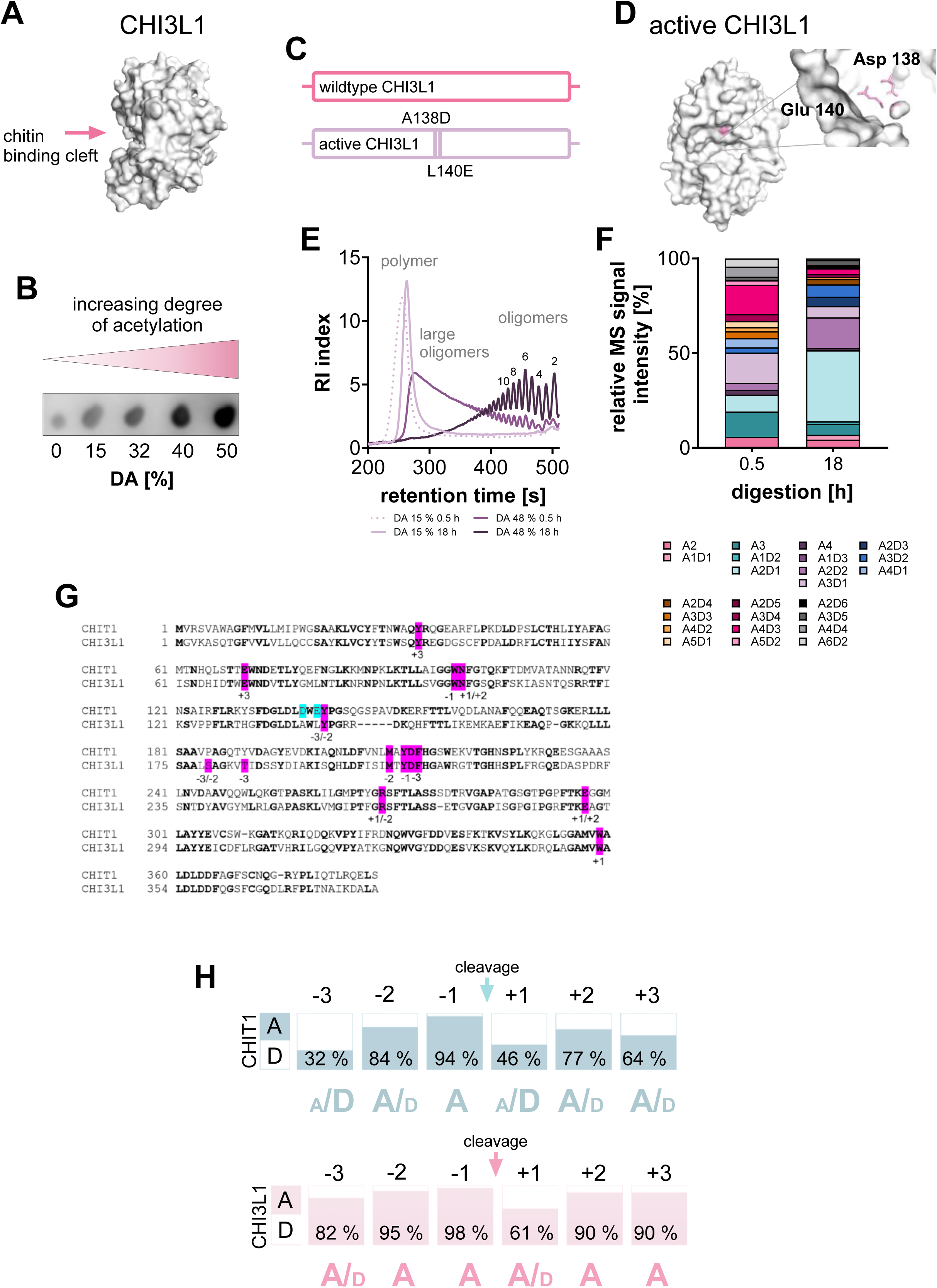
Activated CHI3L1 mutant is able to cleave chitosan and shows a preference for highly acetylated chitosans. A: Structural model of CHI3L1 and its chitin binding cleft. **B:** Binding of CHI3L1 to chitosan with increasing DA, indicating preferential recognition of highly acetylated chitosans. **C:** Schematic comparison of CHI3L1 and activated CHI3L1 mutant with the substitutions A138D and L140E. **D:** Structural model of the activated CHI3L1 variant showing the location of catalytic amino acids in pink. **E:** Size distribution analysis of chitosans (DA 15% & DA 48%) digested with activated CHI3L1 for 0.5 h and 18 h, showing digestion products varying from chitosan polymers over large oligomers to small oligomers. Chitosan with DA = 48% is digested more efficiently. **F:** Product profiles generated by the activated CHI3L1 mutant after incubation with chitosan (DA 48%) for 0.5 h and 18 h, revealing cleavage preferences across defined COS. **G:** Sequence alignment of CHIT1 and CHI3L1 amino acid sequence. Similar amino acids are shown in bold and amino acids related to subsites −3 to +3 are highlighted in pink. The catalytic region is highlighted in cyan. **H:** Subsite preference analysis demonstrating ligand preferences (GlcNAc = A or GlcN = D) of CHI3L1 and CHIT1 across the chitin binding cleft. Activated CHI3L1 shows a stronger preference for acetylated units, especially in the −1 and −2 position. In total, activated CHI3L1 presents more subsites with a high preference for GlcNAc, than CHIT1. Data are representative of n=3.

### Positional screening and molecular simulations identify subsite-1 as the key driver of CHI3L1 binding specificity

To test whether these subsite preferences also apply to enzymatically inactive wild-type CHI3L1, MST binding experiments were performed (**Table 2**, **Supplementary Figure 2**). We analyzed the impact of deacetylation within the binding cleft by testing a panel of chitin hexamers, each containing a single deacetylated GlcN unit at a defined position. The binding affinity for the fully acetylated control (AAAAAA) was K_D_ = 3.2 μM, which is in good agreement with literature data (K_D_ = 6.7 μM).^26^ As a general trend, this positional scan revealed that the location of the GlcN unit is highly critical for the interaction: deacetylation at distal positions was well tolerated, whereas modifications within the interior of the carbohydrate chain significantly impaired binding. Interestingly, subsite +1 emerges as an exception to this trend: introducing a GlcN unit at this internal position (AAAADA) was highly tolerated and caused only a minor decrease in affinity (K_D_ = 11.3 µM). This high tolerance at the +1 subsite directly mirrors our data from the active CHI3L1 subsite preference assay, which showed a lower specificity for acetylation at this position (61%, **Figure 1H**). In contrast, introducing a GlcN unit at the −1 subsite (AAADAA) led to a severe reduction in binding, causing an 80-fold decrease in affinity (K_D_ = 248 µM). This severe loss of affinity perfectly aligns with the strict GlcNAc preference observed for subsite −1 in our active CHI3L1 assay (98%, **Figure 1H**).

**Table 2.**
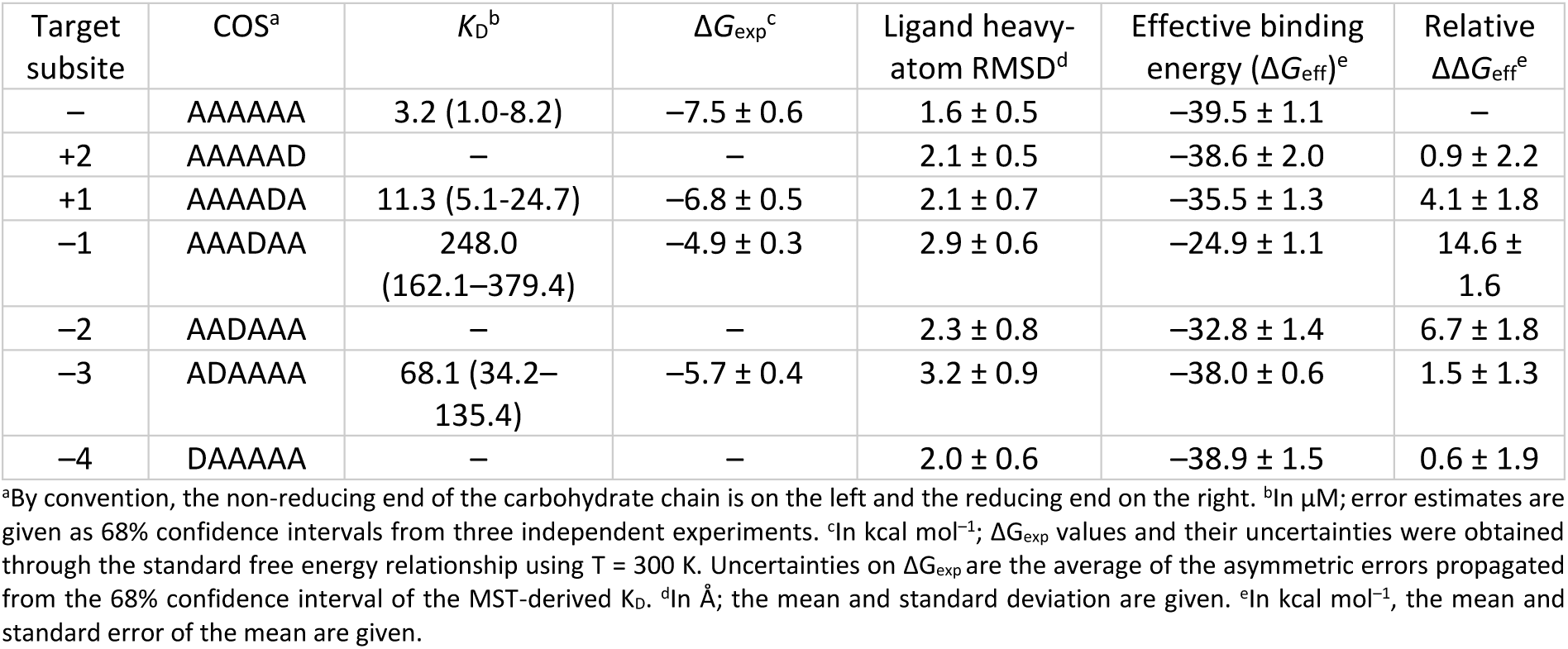
Effect of the Pattern of Acetylation on COS Binding to CHI3L1.

To complement our subsite preference and MST data, all-atom unbiased molecular dynamics (MD) simulations of CHI3L1 were performed using the same panel of mono-deacetylated hexamers. In all simulations the hexamer occupied the subsites –4 to +2, according to the crystal structure (PDB ID: 1HJW)^26^, while the CHI3L1 backbone remained structurally invariant (root-mean-square deviation (RMSD) ∼1 Å) (**Supplementary Figure 2**). To evaluate how the acetylation pattern influences binding, we quantified the structural deviation from the native binding pose and the effective binding energy (ΔG_eff_) for each ligand (**Figures 2A, Supplementary Figure 2**).

**Fig. 2:**
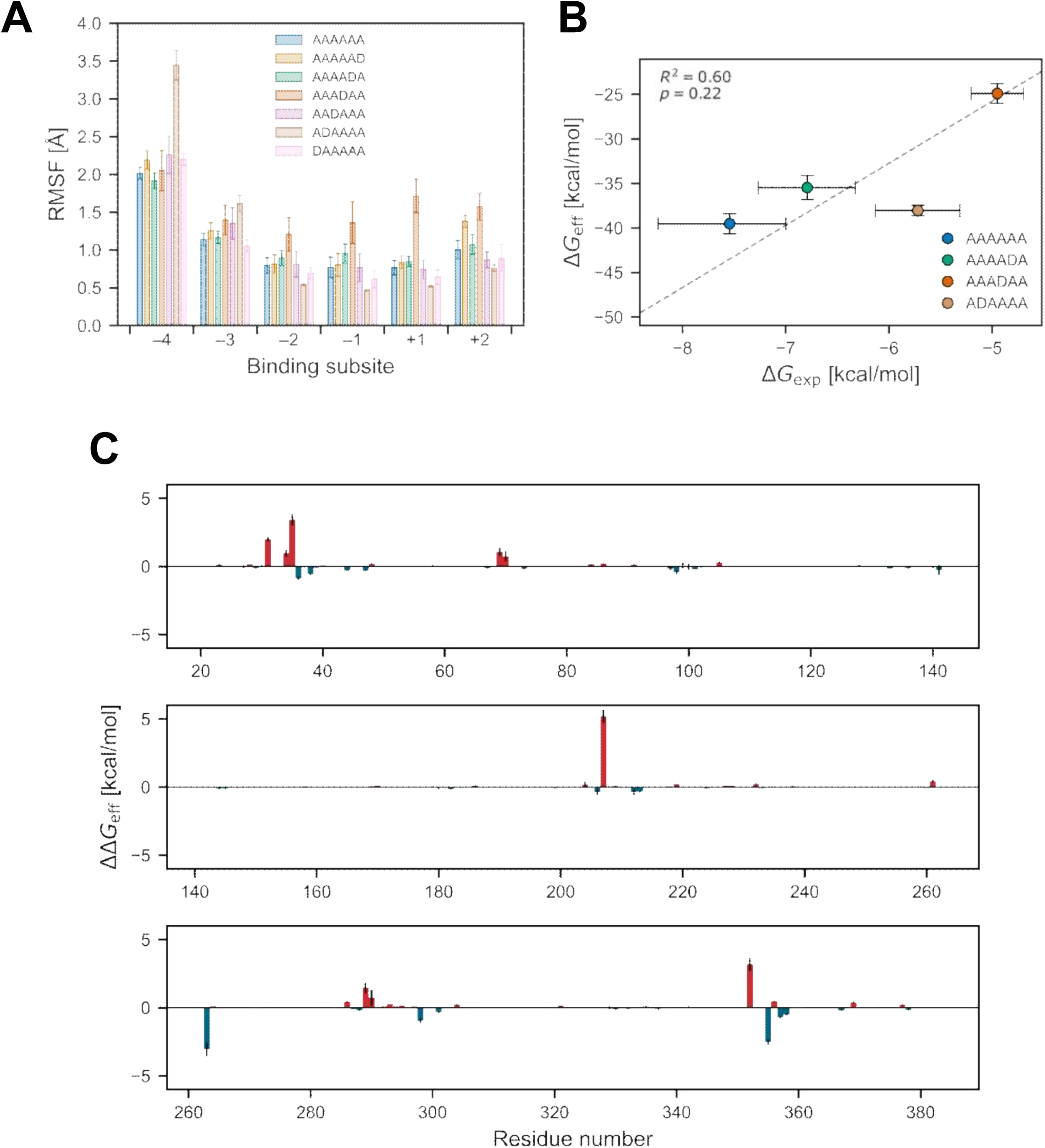
Positional screening and molecular simulations identify subsite-1 as the key driver of CHI3L1 binding specificity. A: Ligand mobility in CHI3L1/COS complexes. RMSF of the different ligands as function of the ligand binding subsite (–4 represents the non-reducing end and +2 the reducing end) calculated using the heavy atoms of each sugar unit. Error bars represent the SEM. **B:** Binding affinity comparison for CHI3L1/COS complexes. Correlation of experimental affinities against MM-PBSA effective binding energies. Error bars denote the SEM for ΔG_eff_ across 12 independent replicas. Experimental free energies (ΔG_exp_) were converted from K_D_ values obtained from MST measurements; error bars represent the 68% confidence interval, derived by propagating the upper and lower K_D_ bound through the standard free energy relationship using T = 300 K. The correlation line is shown in gray. **C:** Per-residue contributions to the relative effective binding energy (ΔΔG_eff_ (AAADAA vs. AAAAAA); GlcN at subsite –1) as calculated by MM-PBSA decomposition. The SEM of ΔΔG_eff_ for each residue was calculated via error propagation (see Materials and Methods).

The simulations revealed that deacetylation within the core subsites (–2 to +1) forced the ligands to displace from their initial positions, causing a loss of crucial interactions and resulting in significantly less favorable binding energies (**Supplementary Figure 2**). This is clearly reflected in the relative binding energies (ΔΔG_eff,_ see Materials and Methods), which underline an highly unfavorable binding for AAADAA (ΔΔG_eff_ = 14.6 ± 1.6 kcal mol^-1^), followed by AADAAA and AAAADA, highlighting the preference of an acetylated unit at these subsites (**Figure 2A**). This trend was further confirmed by analyzing the root-mean-square fluctuations (RMSF) of the ligands: introducing a GlcN unit at subsite –1 (AAADAA) led to markedly increased ligand mobility within the –1/+1 region, explaining the unfavorable shift in binding energy (**Figure 2B**). Interestingly, removing the acetyl group at the distal – 3 subsite (ADAAAA) induced the most significant conformational shift (**Table 2**, **Supplementary Figure 2**), causing the neighboring sugar (subsite –4) to undergo transient unbinding and binding (Figure 1A). However, this rearrangement showed only minor changes in the relative binding energy (ΔΔG_eff_ =1.5 ± 1.3 kcal mol^-1^), suggesting that the ligand adopts an alternative, energetically stable binding pose. Overall, these simulation trends correlate well with our MST experiments (**Table 2**) and subsite preferences data (**Figure 1H**).

To unravel why subsite –1 is the most sensitive subsite for acetyl groups, a per-residue relative binding energy decomposition (ΔΔG_eff_ (AAADAA vs. AAAAAA)) was performed to identify the specific amino acids responsible for this specificity (**Figure 2C**).^65^ This analysis revealed that the affinity loss for AAADAA is primarily driven by unfavorable energetic contributions from three key residues: D207 (ΔΔG_eff_ = 5.2 ± 0.5 kcal mol^-1^), R35 (ΔΔG_eff_ = 3.4 ± 0.4 kcal mol^-1^), and W352 (ΔΔG_eff_ = 3.2 ± 0.5 kcal mol^-1^). Mechanistically, removing the acetyl group at the –1 position disrupts the native, highly favorable CH−π stacking interaction between the sugar ring and the aromatic side chain of W352.^26^ This loss triggers a structural rearrangement of the ligand, forcing the surrounding charged protein residues to incur severe polar desolvation penalties that can no longer be compensated by favorable ligand contacts. In conclusion, these data show a high stringency for specific acetylation patterns, with subsite –1 being responsible for complex stability. Together, the measurements from MST and MD simulations confirm that even a single deacetylation event at this critical position is sufficient to disrupt the energetic network required for high-affinity binding.

### CHI3L1 subsite preferences promote the binding to regularly patterned chitosan

While MST experiments and MD simulations showed how hexameric chitin oligosaccharides (COS) are recognized within the chitin binding cleft, the impact of the chain length on CHI3L1 binding remained to be defined. Moreover and for control experiments, we generated a CHI3L1 mutant lacking the chitin-binding domain (CBmut CHI3L1: CHI3L1 W31A/W99A/W352A) (**Figure 3A**).^26^ First, we analyzed the interaction of CHI3L1 with a series of COS (DP 5–8) using GCI. A channel coated with CBmut CHI3L1 was used as baseline reference (**Figure 3B, Supplementary Figure 3A**). The apparent dissociation constants (K_D_) decreased with increasing DP (**Figure 3B**). To better illustrate how DP affects K_D_, we calculated the impact of the DP change on the apparent K_D_. The analysis revealed that the change of the K_D_ was strongest when changing from DP7 to DP8 (DP7→DP8)(**Figure 3C**), consistent with previous structural data showing that up to eight sugar residues can be accommodated within the CHI3L1 binding cleft (**Figure 3D**).^26^ Static light scattering revealed that COS with DP9 and DP10 form aggregates in aqueous solution (**Supplementary Figure 3B**), preventing their application in our experiments. In further experiments, we used the GCI-derived K_D_ of the DP8 chitin and estimated the total binding energy per cleft (ΔG(cleft, chitin) = −7.76 ± 0.29 kcal mol^-1^) and per residue (ΔG(GlcNAc) = −0.97 ± 0.036 kcal mol^-1^), which is consistent with the calculated energies derived from MST experiments. To estimate the average binding energy of GlcN, we aimed to measure the binding of CHI3L1 to chitosan polymers using fluorescence spectroscopy. To validate this method, we measured again the binding of CHI3L1 to the chitin hexamer (K_D_ = 2.12 ± 0.37 µM, **Figure 3E**). The obtained results were in good agreement with those obtained by GCI and MST confirming the reliability of the fluorescence based assay. Next we measured the binding of CHI3L1 to a series of chitosan polymers with DAs ranging from 0% to 50% (**Figure 3F, Supplementary Figure 3C**) and determined the corresponding K_D_ values. Because each polymer chain contains multiple potential CHI3L1 binding sites, we plotted the percentage of bound CHI3L1 against the concentration of available binding sites. The number of binding sites was calculated as binding sites=c_chitosan_⋅DP/s, where c_chitosan_ is the chitosan concentration and s the number of subsites (here, s=8).

**Fig. 3:**
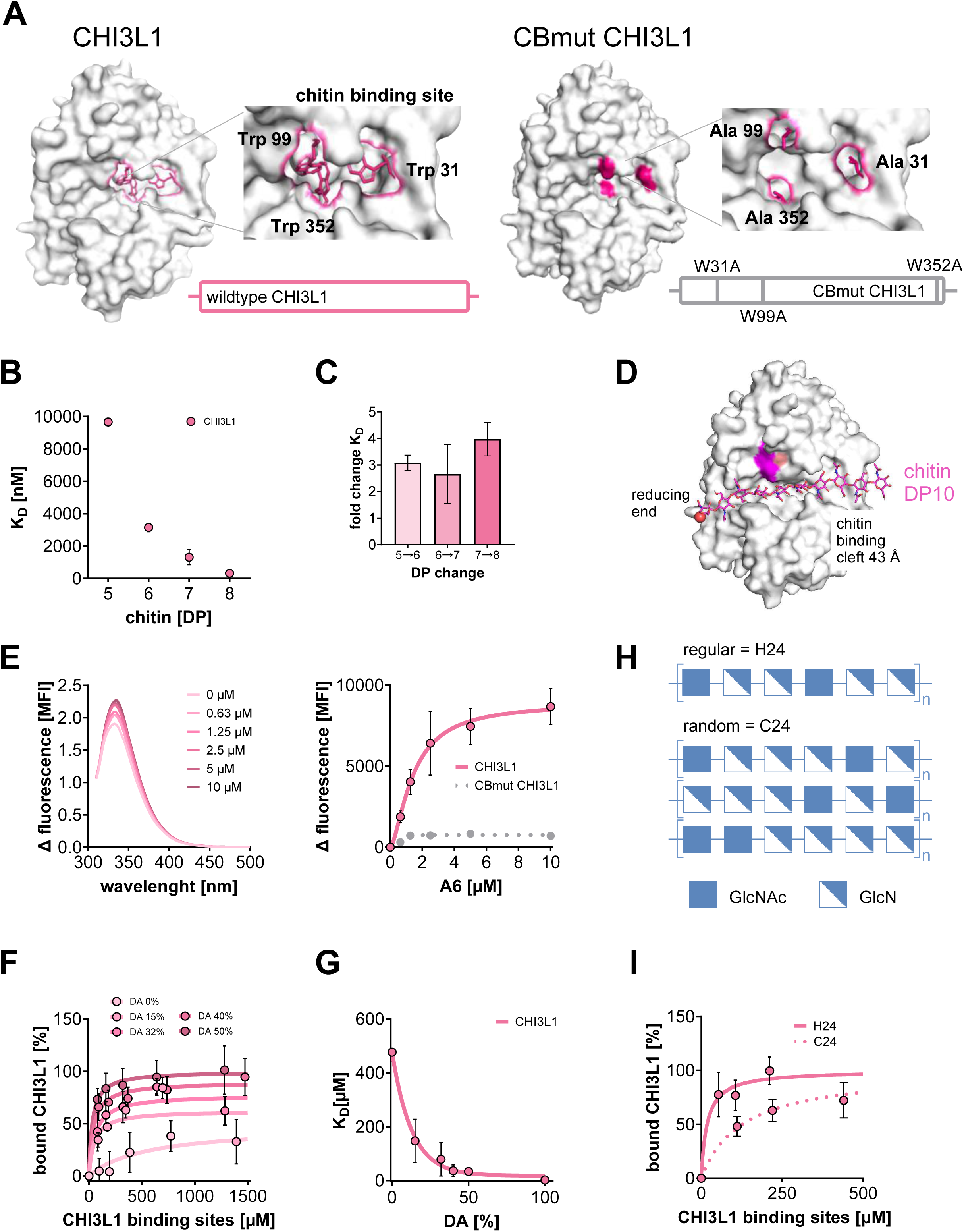
CHI3L1 recognizes chitin and chitosan preferentially binds regular-patterned chitosan. A: Left:structural model of CHI3L1 and its chitin binding cleft. Shown are the three tryptophans (Trp 31, Trp 99 and Trp 352) postulated to mainly contribute to chitin binding in pink. **Right:** structural model of chitin binding deficient mutant (CBmut CHI3L1; W31A, W99A, W352A) and its mutated chitin binding cleft, in comparison to CHI3L1. **B:** Binding affinity of CHI3L1 to chitin oligomers of increasing degree of polymerization (DP 5 - 10), showing enhanced binding (decrease of the apparent dissociation constant K_D_) with increasing chain length as measured by Grating-Coupled Interferometry (GCI). **C:** Impact of the indicated DP changes on the fold-changes of the K_D_. Highest fold change was measured for DP7 → DP8 suggesting 8 subsites in the chitin binding cleft. **D:** Schematic representation of the CHI3L1 binding cleft (43 Å) assuming 8 chitin-binding subsites. **E:** Intrinsic Trp-fluorescence spectra of CHI3L1 interacting with chitin hexamer (A6) at different concentrations (**left**). CHI3L1 was excited with UV light (290 nm). Shown are the emission spectra from 300 to 500 nm (**left**) or the selected emission at 340 nm (**right**). CHI3L1 binds A6 in a concentration-dependent manner as indicated by an increased Δ fluorescence. CBmut CHI3L1 shows no binding to A6, demonstrating the successful mutation of the chitin binding site (**right**). Lines represent fits to a one-site specific binding model. **F:** Change of the intrinsic Trp-fluorescence of CHI3L1 by chitosans with increasing DA (0 – 50%), revealed stronger binding to high acetylated chitosans. Lines represent fits to a one-site specific binding model. **G:** Apparent K_D_ derived from **F** as a function of DA, showing increased affinity with increasing DA. Line represents an empirical exponential one-phase decay fit of the data. **H:** Schematic representation of regularly (H24) and randomly (C24) acetylated chitosan, both with DA = 24%, used for further analysis. Every third unit of a regular chitosan polymer has an increased probability to be a non-acetylated GlcN (D) residue. **I:** Binding of CHI3L1 to H24 and C24, demonstrating stronger binding to the regularly patterned chitosan. Both chitosans had a DA of 24%. Lines represent simulations assuming a one site-specific binding model and the subsite preference of CHI3L1. Data represent mean ± SD, n = 3-10.

While CBmut CHI3L1 showed no detectable binding in control experiments (**Figure 3E**), the K_D_ of CHI3L1 decreased with increasing DA (**Figure 3G**). As expected, the binding to chitosan DA 0% was lowest and close to the detection limit. An empirical exponential decay model was used to correlate DA and K_D_, combining data from chitin hexamer and chitosan polymers. The polymers had a comparable DP of ∼1000 and were therefore much larger than the hexamer. Although the obtained K_D_ values aligned well with the DA, we cannot fully exclude that the differences in DP influenced the measurement. Next, we measured the affinities of CHI3L1 to two chitosans with similar DA (∼24%) but distinct PAs. Both chitosans have previously been characterized: chitosan C24 exhibits a random PA and was produced by chemical *N*-acetylation, whereas chitosan H24, produced by heterogeneous deacetylation, displays a more regular PA characterized by a high frequency of alternating GlcNAc–GlcN–GlcN motifs (**Figure 3H, Supplementary Figure 3C**).^10,11^ Fluorescence spectroscopy revealed a higher affinity of CHI3L1 for H24 than for C24, indicating a preference for a more regularly patterned chitosan (**Figure 3I**). Finally, combining the experimentally determined subsite preferences of the active CHI3L1 A138D/L140E and average binding energies to GlcNAc and GlcN of the CHI3L1, we simulated the interaction of CHI3L1 with both chitosans while explicitly accounting for their distinct PAs (lines, **Figure 3I**). Simulations were performed using a series of chitosan octamers with either random or regular PA patterns (e.g., ADDADDAD). Residue-specific binding energies were derived from the aforementioned average ΔG values and corrected for the respective subsite preferences. The excellent agreement between the simulated (dashed lines, **Figure 3I**) and experimentally measured binding data (dots, **Figure 3I**) demonstrates that the subsite preferences determined for active CHI3L1 A138D/L140E are also valid for CHI3L1. Consistently, CHI3L1 binds with higher affinity to chitosans with a more regular PA than to those with a random PA.

### CHI3L1 enhances TLR2-mediated cell response towards chitosan

In further experiments, we aimed to translate the relevance of the binding between chitosan and CHI3L1 for TLR2-mediated cell activation by employing a reporter cell line that overexpresses human TLR2 (**Figure 4A**). Upon ligand binding, TLR2 signaling promotes the translocation of NF-κB into the nucleus, resulting in the expression of secreted embryonic alkaline phosphatase (SEAP) and thus blue coloration of the culture medium.

**Fig. 4:**
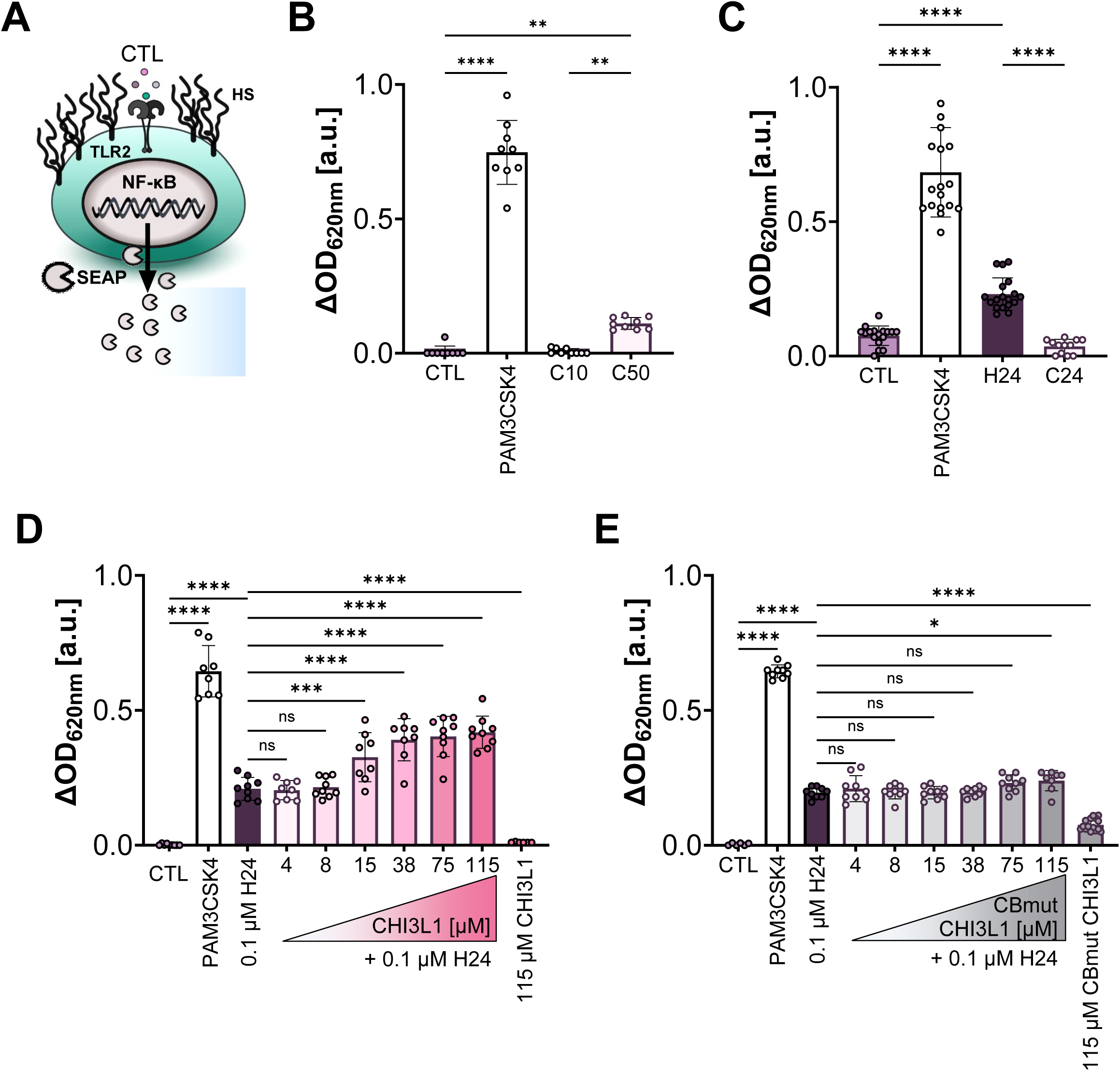
CHI3L1 enhances TLR2 signaling induced by regular patterned chitosan. A: Schematic overview of the HEK-Blue™ hTLR2 reporter cell. TLR2 activation was quantified through a SEAP reporter activity assay recording absorbance at 620 nm. **B:** HEK-Blue™ hTLR2 cells stimulated with chitosan, C10 (DA 10%) and C50 (DA 50%), showing responses to C50. Treatment with C10 led to no activation. PAM3CSK4: positive control. CTL: medium control. **C:** Comparison of regular-acetylation (H24) and random-acetylation (C24) chitosans in the HEK-Blue™ hTLR2 reporter cells, providing a response for H24, but not for C24. **D:** HEK-Blue™ hTLR2 cells stimulated with H24 alone or in combination with increasing concentrations of CHI3L1, demonstrating that CHI3L1 enhanced the cellular response to regular chitosan. **E:** HEK-Blue™ hTLR2 cells stimulated with H24 alone or in combination with increasing concentrations of CBmut CHI3L1. In contrast to CHI3L1, the CBmut CHI3L1 mutant failed to enhance signaling. Data represent mean ± SD, n = 3-5. Statistical analysis was performed using one-way ANOVA with Tukey’ multiple-comparison test, * P<0.05; ** P <0.01; *** P< 0.001; **** P <0.0001.

Chitosans with a more regular PA (H24) are more potent in inducing a TLR2-mediated cell response than randomly patterned chitosans (C24) (**Figure 4B** and **C**).^10^ Interestingly, the response to H24 was even more pronounced than that to the higher acetylated DA 50% chitosan (C50), suggesting that the regular acetylation pattern is more relevant than a higher DA. Chitosans employed for cell stimulation were confirmed to be endotoxin-free (**Supplementary Figure 4A**). Moreover, the reporter cell line employed here shows low responsiveness to endotoxin, indicating that the observed cell activation by chitosans is not related to endotoxin contamination (**Supplementary Figure 4B**). Co-stimulation with CHI3L1 further enhanced cellular response to H24 in a dose-dependent manner (**Figure 4D**). Knockout of TLR2 using CRISPR/Cas9 or neutralization with a TLR2 antibody abolished the response to chitosan (**Supplementary Figure 4C and D**). Furthermore, chitin-binding deficient mutant CBmut CHI3L1 did not enhance the response to H24 (**Figure 4E**).

### CHI3L1 contains two HS-binding sites

In the next part of our work, we analyzed the binding of CHI3L1 to heparin/HS. The ability of CHI3L1 to bind heparin has been described previously^26,27^, and a conserved HS-binding site with the sequence RRDK located at positions 144–147 has been proposed.^66^ In addition to this site, a second potential HS-binding site carrying K or R residues at positions 337, 342 and 344 has recently been suggested (**Figure 5A**).^67^

**Fig. 5:**
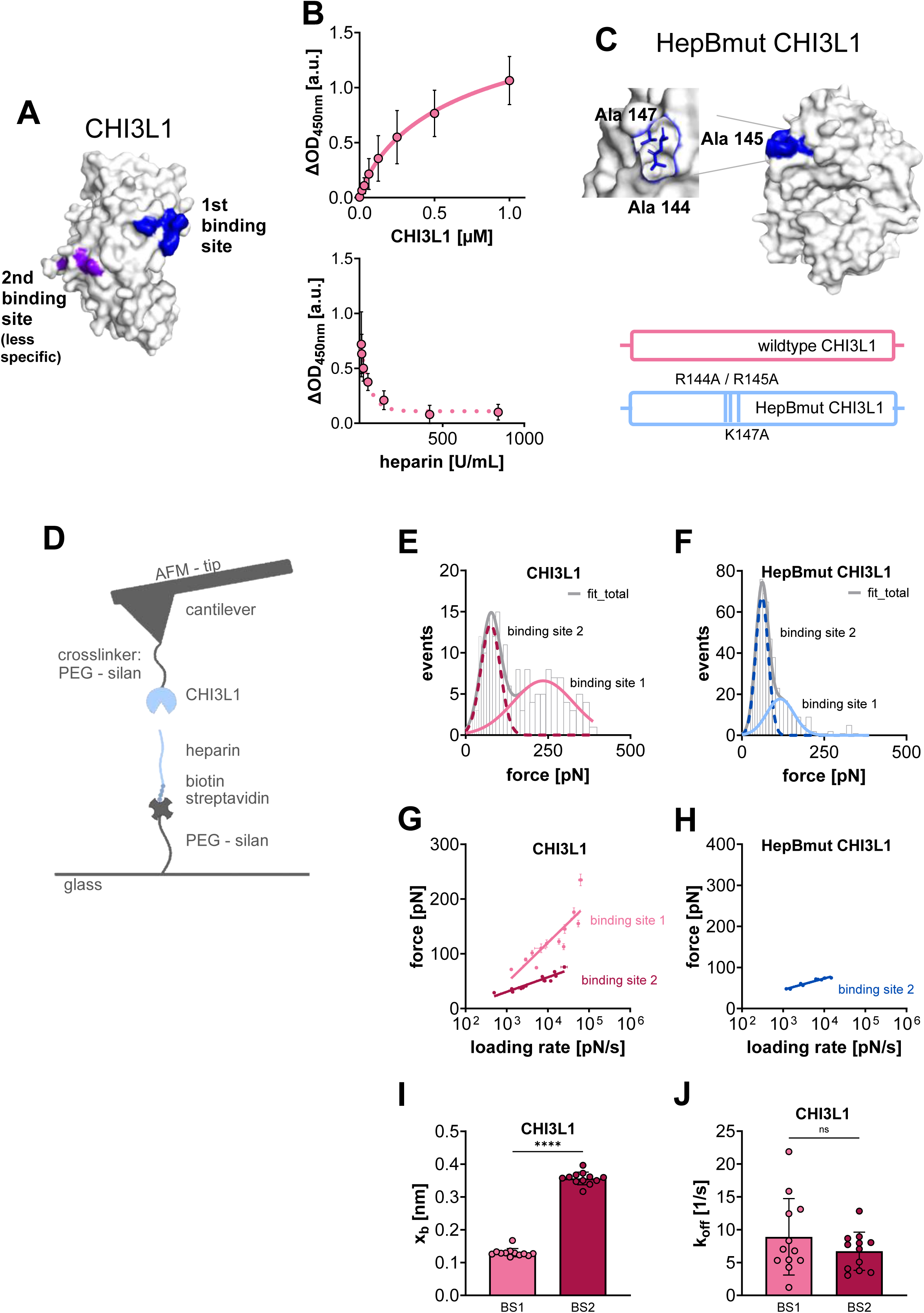
CHI3L1 binds HS motifs through two binding sites. A: Structural model of CHI3L1, indicated are the two postulated HS binding sites. **B:** ELISA like binding assay showing concentration-dependent interaction of CHI3L1 with immobilized unfractionated heparin. Competition with soluble heparin reduces binding to immobilized heparin, indicating displacement of CHI3L1 (K_D_ = 32 ± 29 U/mL). Lines represent fits to a one-site specific binding model. **C:** Structural model of the postulated HS binding site of CHI3L1 and schematic illustration of the HS-binding mutant (HepBmut CHI3L1; R144A, R145A and K147A). Mutated amino acids are shown in blue. **D:** Schematic illustration overview of the atomic force microscopy (AFM)-based single molecule force spectroscopy (SMFS) setup used to analyze CHI3L1-heparin interactions. **E** and **F:** Force-event distribution histograms for CHI3L1 (**E**) and HepBmut CHI3L1 (**F**) interacting with heparin revealing two binding populations (binding site 1 and 2). CHI3L1 exhibits two binding sites. First binding site was diminished in the experiments with the HepBmut CHI3L1 mutant. Gray line represents the sum of two independent Gaussian fits (blue lines). Although the first HS binding site was mutated in the HepBmut CHI3L1 variant, two binding sites were assumed for the data analysis to allow a better comparison to CHI3L1. **G** and **H:** Force - loading rate relationships for CHI3L1 (**G**) and HepBmut CHI3L1 (**H**). **I** and **J:** Barrier widths (*x_b_*) and off rate (*k_off_*) for CHI3L1 obtained from fits of the SMFS data (**G**). Data are shown as mean ± SD from n = 3-4 independent experiments. Statistical analysis was performed using a student’s t test, *** P< 0.001; **** P <0.0001.

Binding of CHI3L1 to surface-immobilized unfractionated heparin (used as functional surrogate for HS) was validated using an ELISA-like binding assay. Addition of soluble heparin competitively reduced the binding of CHI3L1 in a dose-dependent manner (**Figure 5B**). The determined apparent K_D_ was 32 ± 29.13 U/mL, corresponding to ∼11 ± 9.7 µM, indicating that the affinity of CHI3L1 for heparin is of similar magnitude to that for chitin. To characterize the interaction between HS and CHI3L1, we generated a CHI3L1 mutant lacking the 1^st^ HS-binding site (HepBmut CHI3L1: CHI3L1 R144A/R145A/K147A) for control experiments (**Figure 5C**). HepBmut CHI3L1 could be expressed and purified successfully. In contrast, disruption of the 2^nd^ predicted binding site resulted in an unstable protein, which was therefore excluded from analysis. Further, we employed single-molecule force spectroscopy (SMFS). Biotinylated unfractionated heparin was immobilized on a glass coverslip, and CHI3L1 was covalently attached to the tip of an atomic force microscope (AFM) via a polyethylene glycol linker (**Figure 5D**). Binding interactions were probed at different loading rates, and the forces required to rupture individual HS-CHI3L1 bonds were recorded. A representative force histogram revealed a bimodal rupture-force distribution, with one peak centered at 76 ± 29.1 pN and a second peak at 239 ± 109.4 pN (**Figure 5E**). To determine whether these two force populations correspond to the two previously proposed HS-binding sites, we used HepBmut CHI3L1 for further experiments. Before performing SMFS experiments with HepBmut CHI3L1, we confirmed the general integrity of the mutant by assessing its chitin-binding capacity. Because substitution of the three basic amino acids lowers the theoretical isoelectric point (8.7 vs. 8.1), we measured its chitin-binding capacity under different pH conditions (**Supplementary Figure 5B)** and with a series of chitosan polymers with DAs ranging from 0% to 50% (**Supplementary Figure 5C)**. Compared to CHI3L1, we found no detectable differences in chitosan/chitin binding of HepBmut CHI3L1. In SMFS measurements, the force histogram obtained with HepBmut CHI3L1 showed a strong reduction in the high-force peak, whereas the low-force peak remained largely unchanged (**Figure 5F**). This indicates that the high-force population observed for CHI3L1 originates from interactions involving the 1^st^ HS-binding site. To further interpret these findings, rupture forces were plotted as a function of the loading rates (**Figure 5G** and **H**), allowing extraction of kinetic and energetic parameters. **Figure 5I** and **J** show that the low-force peak (2^nd^ binding site) was characterized by a dissociation rate (*k_off_*) of CHI3L1 of approximately 10.1 s⁻¹ and a distance to the transition state (*x_b_*) of 3.7 Å. In contrast, the high-force peak (1^st^ binding site) was characterized by a smaller *x_b_* (0.92 Å) together with a high k_off_ (12 s⁻¹). The small *x_b_* is consistent with the surface-exposed localization (**Figure 5C**) and a very low structural flexibility (**Supplementary Figure 5A**) of the first HS-binding site. The observed high k_off_, together with the relatively large standard deviation may suggest an interaction sensitive to the HS sulfation pattern. Since unfractionated heparin is a heterogeneous mixture, a large fraction of heparin may not bind or only weakly bind to the 1^st^ binding site, resulting in a high standard deviation in the SMFS measurements and a high apparent k_off_. In contrast, the low force interaction between the 2^nd^ binding site and HS suggest a low structural preference but a binding mostly driven by less specific electrostatic interactions.

To test our hypothesis that the 1^st^ binding site preferentially recognizes a distinct sulfation pattern, we analyzed the binding of CHI3L1 and the HepBmut CHI3L1 mutant to 96 defined HS motifs using a glycan microarray (**Figure 6A**). Direct comparison of the two datasets revealed a striking difference in the binding of a specific HS nonamer (9-mer) that was fully *N*- and 6-O-sulfated, contained only a single 2-O-sulfated IdoA residue, and lacked 3-O sulfation. Binding intensities were plotted as a function of the total number of negative charges per HS oligosaccharide (**Figure 6B**). The absolute signal intensities differ between the wild-type and mutant datasets, preventing a direct quantitative scale comparison. Although binding generally correlated with charge density, the 9-mer represented a clear outlier, displaying the strongest binding to CHI3L1. This suggests that CHI3L1 recognizes HS not only through electrostatic interactions, but through specific sulfation patterns. **Supplementary Figure 5D** lists the HS oligosaccharides that showed strong binding to CHI3L1 but markedly reduced binding to HepBmut CHI3L1. Consistent with the strong binding of the 9-mer, other HS oligosaccharides showing relatively strong binding to CHI3L1 also contained *N*- and 6-O sulfations. To better identify potential sulfation patterns preferred by CHI3L1, we clustered the HS oligosaccharides by t-distributed stochastic neighbor embedding (t-SNE) using structural features including the length, number of charge and their pattern of sulfation (**Figure 6C**). After superimposing the clusters with the binding strength of CHI3L1 (**Figure 6D**) and HepBmut CHI3L1 (**Figure 6E**), we found that minimum CHI3L1 binding requires *N*- and 6-O sulfation; however, presence of 2-O sulfation further enhanced CHI3L1 binding. In contrast 3-O sulfation seemed to be not relevant.

**Fig. 6:**
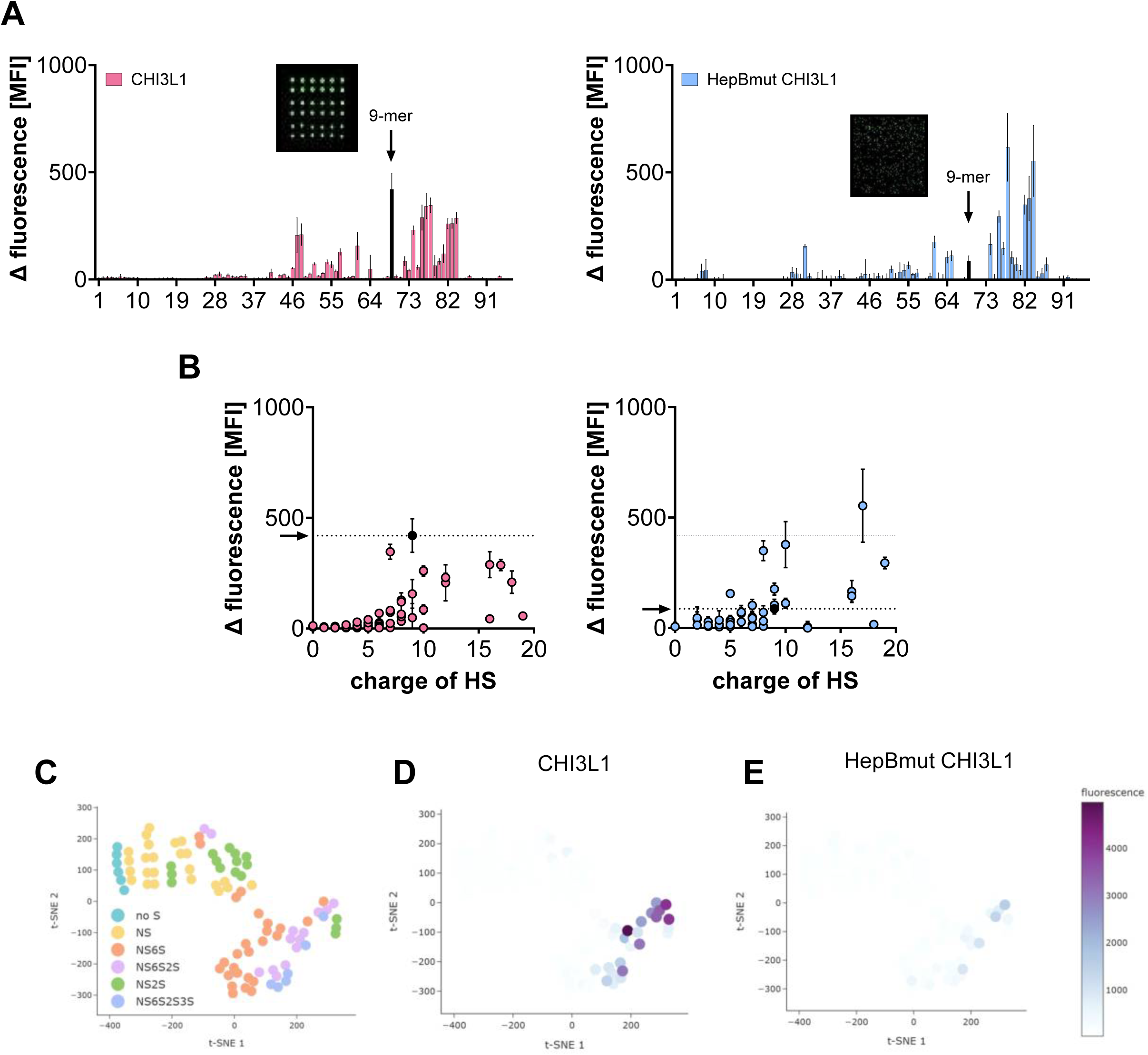
CHI3L1 binds HS motifs sulfation pattern sensitive. A: HS microarray analysis of CHI3L1 (**left**) and HepBmut CHI3L1 (**right**) against 96 different HS motifs. The 2S-6S 9-mer motif (9-mer), found to be strongly bound by CHI3L1 but weakly by HepBmut CHI3L1, is highlighted and used for further analysis. The insets show microarray raw data of the 2S-6S 9-mer. **B:** Plot derived from HS microarray showing the relationship between protein binding intensity and HS charge. CHI3L1 shows increasing binding with higher sulfation levels, whereas the HepBmut CHI3L1 mutant displayed an altered binding profile. **C:** Clustered HS oligosaccharides by t-distributed stochastic neighbor embedding (t-SNE) using structural features including the length, number of charges, and their pattern of sulfation. **D and E:** Superimposing clusters with the binding strength of CHI3L1 (**D**) and HepBmut CHI3L1 (**E**). Minimum CHI3L1 binding requires *N*- and 6-O sulfation; however, the presence of 2-O sulfation further enhanced CHI3L1 binding. In contrast 3-O sulfation seemed to be less relevant.

To further validate the results of both microarrays, we tested the binding of the pentasaccharide fondaparinux and the 9-mer to CHI3L1 and HepBmut CHI3L1 using fluorescence spectroscopy (**Supplementary Figure 5E**). Both protein variants bound fondaparinux with a similar affinity; however, the high standard deviation pointed to a weak and less favored interaction. The 9-mer showed reduced affinity to HepBmut CHI3L1, thus consistent with the microarray data. However, the change in intrinsic fluorescence of CHI3L1 upon binding to the HS oligosaccharides was very small and, compared to the aforementioned experiment with chitosans, close to the detection limit of the method. Therefore, we confirmed the site-specific interaction of CHI3L1 with HS using quartz crystal microbalance with dissipation monitoring (QCM-D).

### The first HS-binding site of CHI3L1 prefers 6-O-sulfated HS

To further characterize properties of the interaction, including its stability, specificity and the ability of CHI3L1 to bind multiple HS, we conducted QCM-D analysis with dissipation monitoring (**Figure 7A**).^60^ QCM-D is based on an oscillating quartz crystal sensor disc, whose frequency is related to its mass. When molecules interact with the sensor surface, its mass increases, which is monitored as a real-time decrease in frequency (ΔF). We functionalized the sensor as described in **Supplementary Figure 6B** and coated it with biotinylated heparin to mimic the cell surface. The chains are free to move laterally on the sensor surface. The second parameter measured is energy dissipation (ΔD), which reflects the viscoelastic properties of the surface layer. Low dissipation indicates a rigid, stiff layer (crosslinking), whereas high dissipation corresponds to a soft, hydrated, and more flexible layer (**Supplementary Figure 5F**).

**Fig. 7:**
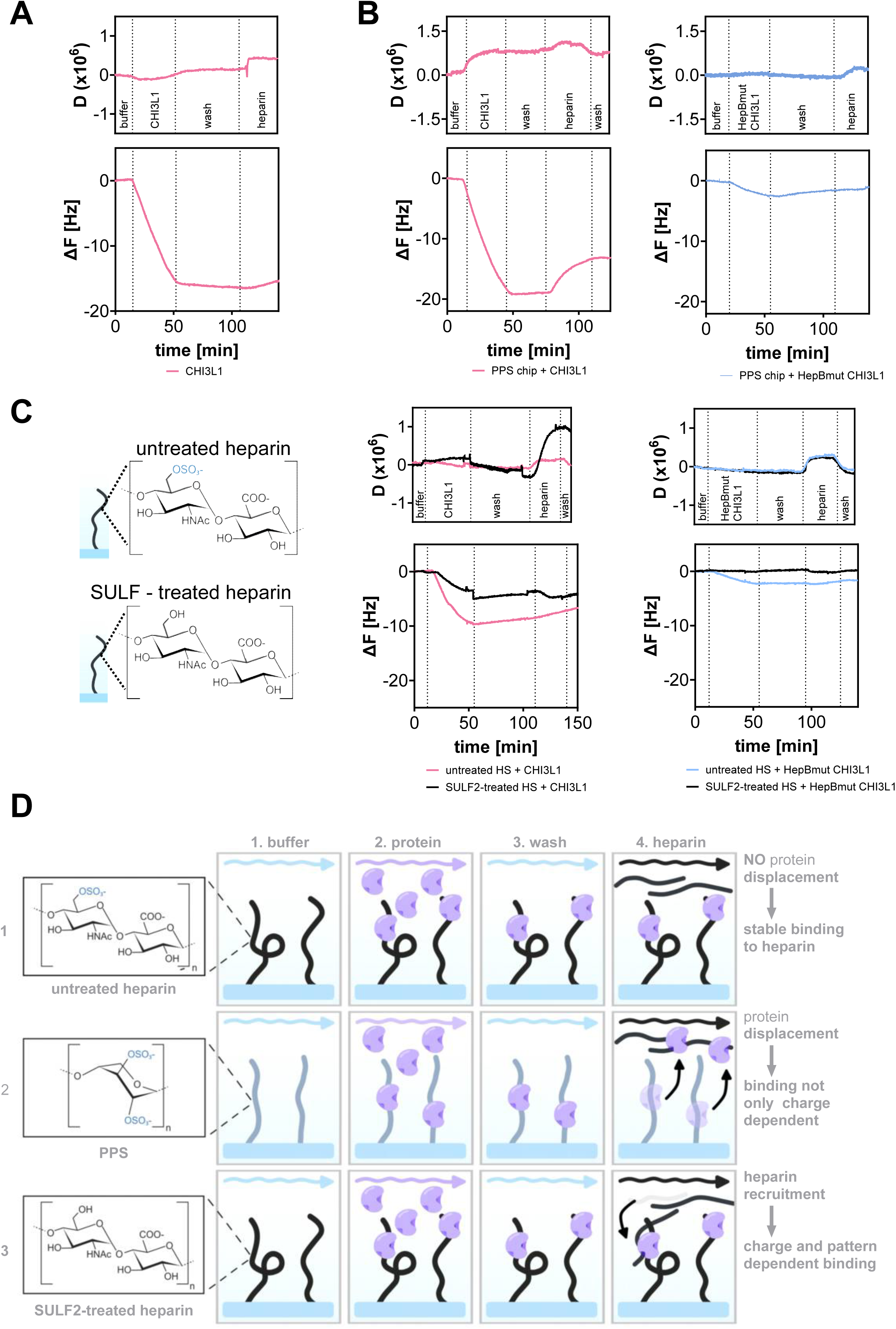
Two binding sites mediate sulfation-dependent CHI3L1 HS / heparin interactions. A: QCM-D analysis showing CHI3L1 binding to immobilized heparin on the chip surface. Protein binding results in decreased resonance frequency (ΔF) (mass adsorption). Followed by slight increase in dissipation (D), reflecting weak HS chain disentanglement. **B:** QCM-D analysis of CHI3L1 and HepBmut CHI3L1 binding to PPS coated chip surfaces. CHI3L1 bound to PPS as indicated by decreasing ΔF. ΔF increased upon addition of soluble heparin, indicating dissociation of surface-bound CHI3L1. HepBmut CHI3L1 showed weaker binding compared to CHI3L1 and dissociated continuously during the washing and heparin perfusion phase. **C:** Comparison of CHI3L1 and HepBmut CHI3L1 binding to untreated and SULF-treated heparin surfaces by QCM-D. **Left**: Schematic overview of the chemical structure of SULF-treated and untreated heparin. **Middle**: Binding of CHI3L1 to SULF-treated and untreated heparin was of a comparable magnitude (ΔF). Washing of the sensor in the 3. phase of the experiment slightly displaced CHI3L1 from the SULF-treated heparin but not from the untreated heparin. Perfusion of the sensor chips with heparin (4. phase of the experiment) resulted in a recruitment of the soluble heparin to the SULF-treated sensor. Heparin was not recruited to the untreated sensor. Increase in dissipation (D) reflected HS chain disentanglement on both sensors. **Right:** HepBmut CHI3L1 showed weaker binding compared to CHI3L1 (**middle**) on the untreated chip. Binding to the SULF-treated chip was nearly abolished. **D:** Schematic overview of different types of binding to immobilized heparin/PPS. CHI3L1 binding to untreated heparin (1), to PPS (2) and to SULF2-treated heparin (3).

In the first experiment (**Figure 7A**) we measured the interaction between CHI3L1 with heparin. CHI3L1 bound to the heparin-coated surface as indicated by decrease in frequency (-ΔF). After binding, the signal remained stable during washing (no change in ΔF), indicating stable interactions. In contrast to thrombin, a prototypic nonspecific HS-binding protein with one HS interaction site,^68^ which dissociates from the surface during the washing step (**Supplementary Figure 5F**). We observed no substantial change in dissipation, indicating no significant crosslinking or bridging of the heparin chains. When we injected soluble heparin for competition, most surface-bound CHI3L1 remained attached, indicating stable binding of CHI3L1 to immobilized heparin.

Performing the same experiment with HepBmut CHI3L1 (lacking the first HS binding site) revealed reduced binding. This indicates that the 2^nd^ binding site contributed only marginally to HS binding.

To exclude the possibility that this interaction is purely electrostatic, we performed experiments using pentosan polysulfate (PPS)-functionalized chips instead of heparin. PPS is a highly and uniformly sulfated polysaccharide, with higher negative charge than heparin (three sulfo groups per disaccharide, compared to 2.7 sulfo groups per disaccharide).^69^ **Figure 7B** shows that CHI3L1 bonds to PPS similarly as heparin. However, perfusion with soluble heparin displaced CHI3L1, indicating CHI3L1 preferentially binds structural features of heparin rather than to high negative charge alone.

The binding of HepBmut CHI3L1 to PPS was comparably weak, indicating that an increased negative charge cannot compensate for the loss of the 1^st^ HS binding site.

Our microarray data suggested that 6-O sulfation is strongly preferred by the 1^st^ binding site. Therefore, we performed the QCM-D experiment in parallel using untreated and SULF2-treated heparin lacking 6-O sulfation on the chip surface (**Figure 7C**). CHI3L1 bound significantly less to SULF2-treated heparin. However, upon injection of soluble heparin, we measured a decrease in ΔF and an increase in ΔD, indicating that the heparin is deposited onto the surface-bound CHI3L1.

We repeated the experiment with the HepBmut CHI3L1 mutant (**Figure 7C**). We observed that the loss of 6-O sulfation abolished HepBmut CHI3L1 binding completely.

Taken together, these results indicate that the 1^st^ HS-binding site is relevant for the initial binding of CHI3L1 to the surface immobilized heparin. The 2^nd^ HS-binding site is less specific and may only stabilize CHI3L1 binding. The deposition of heparin to the slide coated with heparin lacking 6-O sulfation suggest that one of the two binding sites is occupied once soluble heparin is perfused over the sensor, while the other binding site stays in contact with the surface-immobilized heparin. We postulate that the 1^st^ binding site, which preferentially binds 6-O sulfated heparin, switches to the soluble heparin, while the 2^nd^ binding site keeps its interaction with the heparin at the sensor surface.

Our interpretation of the QCM-D data is summarized in **Figure 7D**.

### CHI3L1 binding to cell surfaces depends on HS

To validate the relevance of HS for the binding of CHI3L1 to cell surfaces, we used previously established CRISPR/Cas9-engineered model cell lines in subsequent experiments.^57,58^ Here, we analyzed cells with *EXT1*, *NDST1* and *HS6ST1* gene knockouts in comparison to CTL cells expressing non-targeting sgRNAs. We have previously shown that EXT1KO cells completely lack HS.^58^ Moreover, we found that HS chains in HS6ST1KO cells were more flexible, whereas the chain length was comparable to HS of CTL cells.^57^ Consistent with our previous findings, size exclusion chromatography suggests only a slight increase in HS chain length upon HS6ST1 gene knockout (**Supplementary Figure 7A**). Likewise, HS6ST1KO did not affect the chondroitin sulfate expression (**Supplementary Figure 7B**). To quantify the HS levels at the cell surface of HS6ST1KO cells, we used flow cytometry and a HS-directed antibody (clone 10E4) (**Figure 8A**). In comparison to control cells (CTL), we found that HS6ST1KO cells showed increased 10E4 antibody binding, most likely due to an increased presence of 10E4 epitopes along the HS chain. In line with previous literature, EXT1KO and NDST1KO abolished 10E4 binding, indicating either complete loss of HS chains or a lack of 10E4 epitopes associated with *N*-sulfated residues^70^. Next, we incubated the cells with CHI3L1 and HepBmut CHI3L1 and quantified the accumulation of the proteins at the cell surface by flow cytometry (**Figure 8B**). Compared to the CTL cells, complete loss of HS (EXT1KO), lack of *N*-sulfation (NDST1KO) and alterations in 6-O sulfation (HS6ST1KO) significantly diminished the binding of CHI3L1. The binding of HepBmut CHI3L1 was low for all tested cell lines, further underlining that the 1^st^ HS binding site of CHI3L1 is sensitive to HS sulfation.

**Fig. 8:**
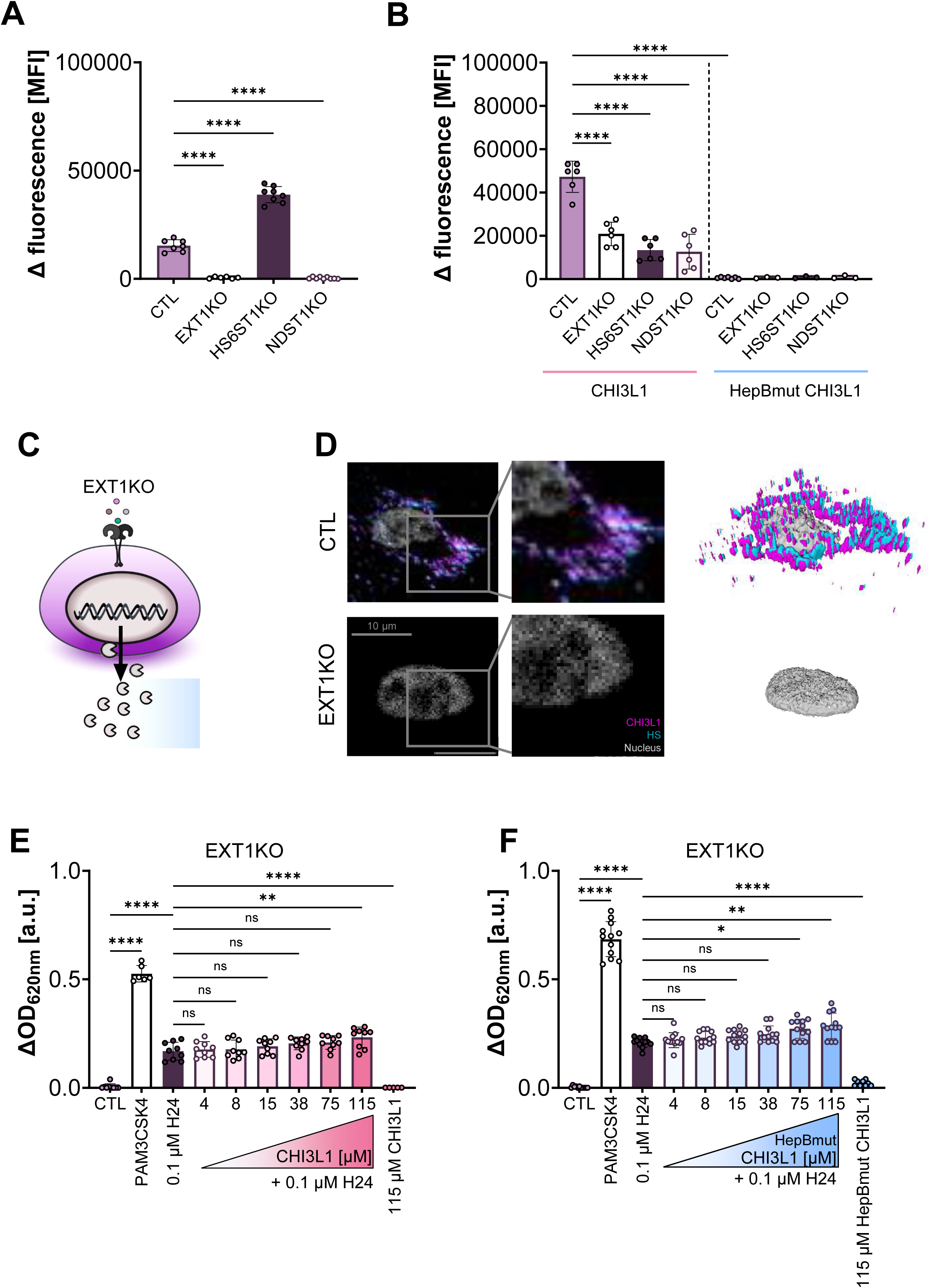
HS determines CHI3L1 binding and downstream cellular responses. A: Flow cytometric analysis of cell surface HS on CTL, EXT1KO, HS6ST1KO and NDST1KO cells. HS was stained by the HS antibody clone 10E4. Binding of 10E4 was abolished in EXT1KO and NDST1KO cells. Enhanced 10E4 binding to HS6ST1KO cells indicate the presence of more antibody binding epitopes. **B:** Flow cytometric analysis of CHI3L1 and HepBmut CHI3L1 binding to cell surface. Loss of HS biosynthesis or altered sulfation reduces binding of CHI3L1. HepBmut CHI3L1 binding to all tested cells was strongly reduced. **C:** Schematic overview of HEK-Blue™ hTLR2 reporter cells incapable of HS biosynthesis (EXT1KO). TLR2 activation was quantified through a SEAP reporter activity assay recording absorbance at 620 nm. **D:** Representative immunofluorescence images (**left**: 2D; **right**: 3D rendered images) captured by structure illumination microscopy (nucleus = grey; CHI3L1 = pink; HS = blue) of CTL and EXT1KO HEK-Blue™ hTLR2 cells. CHI3L1 and HS are co-localized in CTL cells. CHI3L1 binding to EXT1KO cells (no HS expression) was strongly decreased. Scale bar = 10 µm. **E:** EXT1KO HEK-Blue™ hTLR2 cells stimulated with H24 and increasing concentration of CHI3L1. Loss of EXT1 inhibits the CHI3L1-dependent enhancement of signaling, indicating that HS is required for TLR2-mediated cell activation. **F:** HEK-Blue™ hTLR2 cells stimulated with H24 and increasing concentration of HepBmut CHI3L1. Loss of 1^st^ HS binding site inhibits the CHI3L1-dependent enhancement of signaling. Data represent mean ± SD, n = 3-5. Statistical analysis was performed using one-way ANOVA with Tukey’s multiple-comparison test, * P<0.05; ** P <0.01; *** P< 0.001; **** P <0.0001.

Next, we deleted EXT1 in the TLR2 reporter cell line by CRISPR/Cas9 in order to abolish HS biosynthesis (**Figure 8C**). Structured illumination microscopy indicates that in comparison to the control (CTL) cells, knockout of EXT1 abolished HS expression and CHI3L1 binding to the cell surface (**Figure 8D**). On CTL cells, CHI3L1 co-localize with HS. Consistent with the microscopy data, the response of the HS-deficient cells to H24 was not enhanced upon addition of CHI3L1 (**Figure 8E**). Furthermore, absence of the 1^st^ HS binding site markedly reduced the CHI3L1-mediated enhancement of cell activation, comparable to the level of cells lacking HS (**Figure 8F**).

Taken together, our data suggest that CHI3L1 shuttles chitosan to TLR2 at the cell surface, employing HS as a co-receptor. In addition, the ability of CHI3L1 to accumulate chitosan at the cell surface depends on the 1^st^ HS binding site. Our data are schematically summarized in **Figure 9**.

**Fig. 9:**
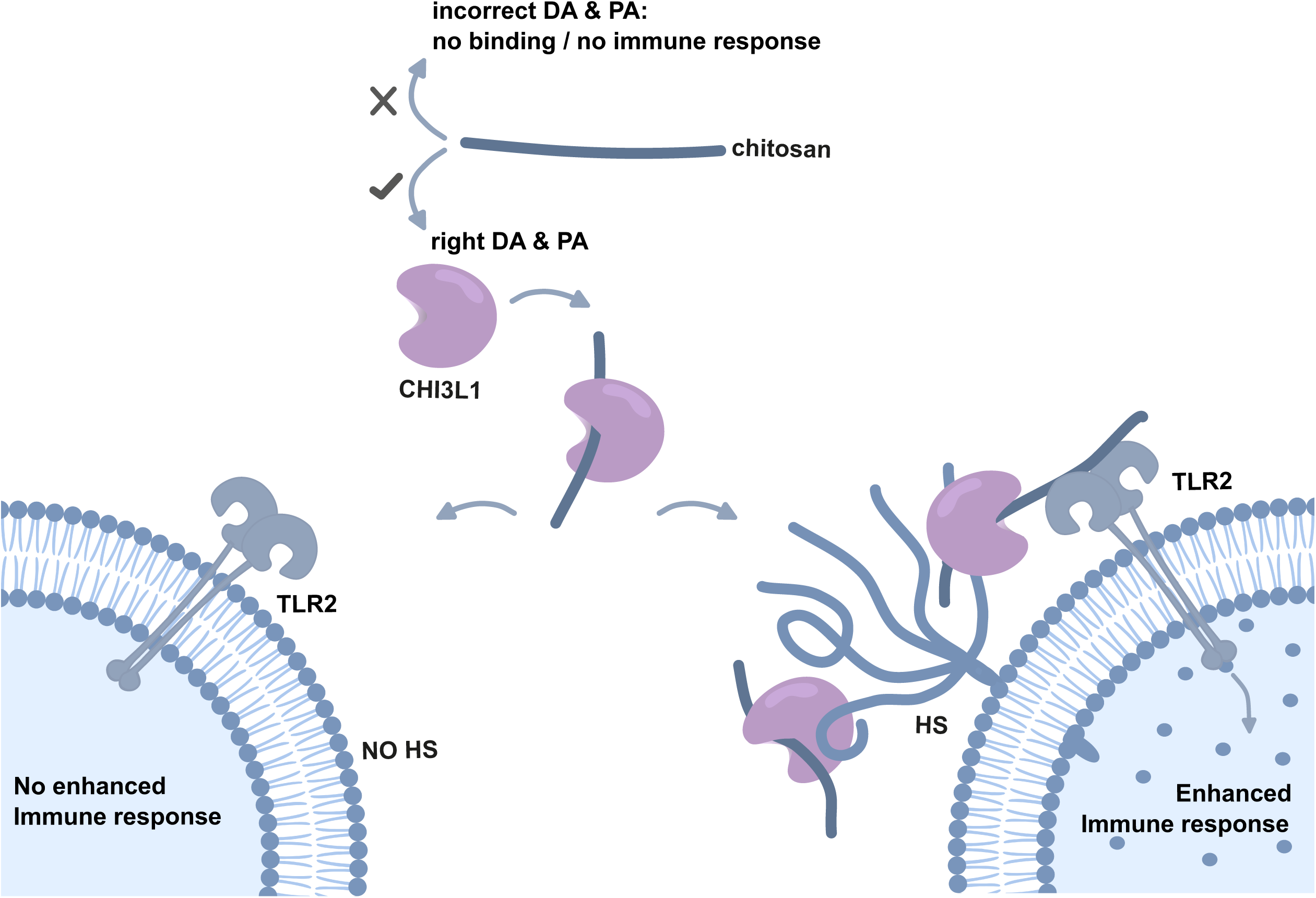
Chitinase-3-like protein 1 promotes chitosan-induced TLR2 signaling through interaction with HS. CHI3L1 recognizes chitosan in a DA- and PA-dependent manner, showing a preference for regularly patterned chitosan. Subsequently, CHI3L1 shuttles the chitosan to the cell surface by interacting with surface-exposed HS. The accumulated chitosan then ligates to TLR2, triggering intracellular signaling. Notably, TLR2 signaling is not activated in the absence of HS or when the DA and PA of chitosan are incorrect.

## Discussion

The application of chitosan in biomedical applications is often connected to its high biodegradability. Enzymatic degradation of chitosan by chitinases such as, CHIT1 depends largely on the DA. Low-acetylated chitosan, which is non-inflammatory, is, however, only weakly degraded by human chitinases. In contrast, highly acetylated chitosan, which is rapidly degraded, is pro-inflammatory.^5,9^ A better understanding of the pro-inflammatory properties of chitosan and the signaling involved is therefore essential to develop tailor-made chitosans with distinct physico-chemical as well as biological properties required for distinct biomedical applications.

Here, we confirmed the previously postulated impact of the PA on the pro-inflammatory properties of chitosans.^10^ While a random PA showed only low TLR2-mediated cell activation, a more regular PA promoted a strong response. Our data further suggest that the enhanced inflammatory response is linked to the subsite preference of CHI3L1. Using GCI, MST, an MS-based workflow and MD simulations, we comprehensively mapped the subsites of CHI3L1. These analyses identify subsite –1 as the central electronic and structural hotspot of the CHI3L1 binding cleft. Removing a single acetyl group at this position induces a severe 80-fold loss in affinity, because it disrupts the native CH–π stacking interaction with W352 and forces the ligand into an unfavorable structural rearrangement. ΔG_eff_ values predicted by effective binding energy computations are low, the magnitudes and fluctuations of these values fall well within the ranges reported in comparable studies.^71^ Based on the detailed characterization of the chitin binding cleft, we provide evidence that the regular arrangement of the acetyl groups enhances the interaction of chitosan with CHI3L1. Together with our finding that CHI3L1 enhances the pro-inflammatory cell activation, we propose that CHI3L1 acts as a shuttle between chitosan and the cell surface. We further showed that CHI3L1 accumulation at the cell surface was mediated by HS, suggesting a triangular relation between CHI3L1, HS and TLR2 that regulate the pro-inflammatory properties of chitosan. Previous physico-chemical studies of chitosan indicate that the acetylation of chitosan significantly affects its solubility, charge distribution, and conformation in solution.^72–74^ Accordingly, low acetylated chitosan (DA < 20%) is a flexible polymer chain with a high positive charge at acidic pH. The presence of negatively charged proteins or other macromolecules may induce secondary aggregation and polyelectrolyte complex formation in biological fluids. Highly acetylated chitosan exhibits only isolated positive charges and polymer chains interact with each other by hydrophobic interactions and hydrogen bonds, promoting the formation of primary aggregates.^73^ To our knowledge, physico-chemical properties of chitosans with a regular PA have not yet been studied. As a result of the regular arrangement of acetyl groups in these chitosans, highly acetylated regions, predominantly characterized by hydrophobic interactions, are likely missing and the formation of primary aggregates might be absent. In addition, the lack of highly positive charges along the chain might also lead to reduced electrostatic interactions with proteins in biological media, suggesting the absence of secondary aggregation as well. Consequently, chitosans with a regular PA might represent a relatively flexible polymer chain with a comparable high bioavailability in biological media, which may contribute to an enhanced pro-inflammatory effect.

Previously, the interaction of CHI3L1 with HS has been studied and connected to angiogenesis in tumors.^75^ In tumors, angiogenesis is controlled by tumor-associated macrophages (TAM). TAMs are known to release CHI3L1 ^76^ and to express TLR2 at considerable levels^77^. Chitin and chitosans are absent in tumors; however, previous MD simulations suggest that CHI3L1 can interact with hyaluronan^66^, which is a major component of the tumor microenvironment^78^. Low molecular weight hyaluronan is known to promote inflammation and angiogenesis in tumors^79,80^ and to be a potent TLR ligand^81^. Further research will be required to reveal the role of CHI3L1 in tumor inflammation and angiogenesis and whether low molecular weight hyaluronan signaling in tumors is supported by CHI3L1. Recently, in a study investigating inflammatory arthritis, conditional SULF2 knockout in macrophages promoted joint inflammation,^82^ suggesting an essential contribution of 6-O sulfated macrophage-associated HS to the development of the disease. Interestingly, CHI3L1 is known to be elevated in patients with arthritis^83^ and to be linked to macrophage infiltration.^84^

Here, we found that CHI3L1 has two HS binding sites. The interaction between the 1^st^ binding site and HS appears to be 6O-sulfation-dependent and, as suggested by glycan microarrays, supported by IduAS2. Additional 3-O sulfation does not appear to enhance this interaction but may even hinder binding to the first site. In contrast, the 2^nd^ binding site seems more permissive and accommodates HS chains with a broader range of sulfation patterns.

While the *k_on_* determined by SMFS suggests comparably strong interactions between HS and both binding sites of CHI3L1, QCM-D measurements revealed that binding of CHI3L1 to surface-immobilized heparin was strongly reduced when the 1^st^ binding site was mutated (HepBmut CHI3L1). Because SMFS experiments provide information only on *k_off_* but not on the corresponding *k_on_*, these data alone cannot fully explain the observed discrepancy. However, the strongly reduced binding of the HepBmut CHI3L1 mutant recorded in the QCM-D experiments suggests that the mutation primarily affects the association rate, suggesting a markedly reduced *k_on_*. This indicates positive cooperativity between both HS binding sites, where the 1^st^ binding site initiates HS binding and the 2^nd^ binding site stabilize the interaction with HS. Previous studies on the morphogen Hedgehog (Hh), which contains two equivalent HS-binding sites, suggested that Hh can move along sulfation gradients.^60^ Consistent with this model, QCM-D analysis showed that Hh remained bound to the sensor surface during washing and even during perfusion with lower-sulfated heparin, whereas perfusion with equally sulfated or more highly sulfated heparin displaced the protein. Comparable experiments with CHI3L1 showed that CHI3L1 remained stably bound to surface-immobilized heparin during washing (probably because of the relatively low k_off_ of the 2^nd^ HS binding site) and only few proteins where displaced from the surface upon injection of heparin of comparable sulfation. In contrast to the observations reported for Hh, perfusion of the sensor chip with more highly sulfated heparins did not displace CHI3L1. Instead, the soluble heparin was recruited to the sensor surface. Based on these findings, we propose a binding model in which two HS binding sites of CHI3L1 exhibit different binding properties. The 1^st^ binding site is more sensitive for 6-O-sulfation and mediates initial interaction with HS, whereas the 2^nd^ binding site contributes to the stability of the HS binding. Recently, a heparin column-based binding assay suggested that the binding of CHI3L1 to heparin is allosterically decreased in the presence of COS.^85^ However, using our heparin ELISA and SMFS setup, we were not able to measure a strong allosteric impact of chitin binding on CHI3L1 (**Supplementary Figure 8**).

Altogether, we found that CHI3L1 cooperates with HS to accumulate chitosan at the cell surface to trigger a TLR2-mediated response. A comparable PAMP collection and enrichment process has previously been shown for the recognition of LPS by TLR4: LPS is captured in the extracellular space by the LPS-binding protein, which interacts via CD14 with the cell surface and presents LPS to TLR4.^86,87^ Lack of the LPS-binding protein drastically decreases the LPS-induced immune reaction in mice.^88^ In line with our findings, CHI3L1 deficiency or antibody-based neutralization significantly reduced inflammation in a hepatitis model.^89^

In conclusion, we provide evidence that chitosan is recognized by CHI3L1 in a DA- and PA-dependent manner, promoting a pro-inflammatory response involving HS and TLR2 signaling. While such inflammatory properties are generally considered undesirable for biomedical applications such as tissue regeneration, they may be advantageous in the context of vaccination, where chitosan could enhance immune responses to vaccine antigens. Although chitosan-based nanoparticles have previously been explored for vaccine delivery, the deliberate use of structurally defined pro-inflammatory chitosan as an adjuvant has, to our knowledge, not yet been investigated. Moreover, our findings provide further evidence that low-acetylated chitosan is immunologically inert and thus useful for a variety of biomedical applications such as tissue regeneration.

## Authors’ contributions

A.G. and C.G. conceived and designed the study. C.G. supervised the project, and C.G., J.C., H.G., B.M.M, H.M. acquired funding. A.G. and C.G. wrote the manuscript. A.G. performed the majority of the experiments and analyzed the data. N.R. and Ö.K. performed molecular simulations and MST experiments, resp., analyzed the data. S.N. performed flow cytometry experiments and analyzed the data, C.I. performed fluorescence microscopy, M.J.H. and S.C.L. performed SEC-RI-MS measurements and subsite preference analysis, J.F. performed and analyzed QCM-D measurements, E.E.A. performed and analyzed GCI measurements, A.N.W., T.T., F.Y., E.G., G.R. provide TLR2 reporter cells, chitosans, COS, heparin or Sulfs. M.K.G. performed HS chain length analysis. S.N., C.I., M.J.H., J.F., E.E.A., S.C.L., N.R., Ö.K., M.D., J.C., H.G., R.R.V., A.N.W., H.M., A.OM., K.G., and B.M.M. discussed the data and edited the manuscript. All authors reviewed and approved the final manuscript.

## Acknowledgments

This study was funded by the German Research Foundation within the Priority program Program 2416 CodeChi GO 2528/10-1 (525894428), OS 497/7-1, 525934838, 524352358, GO 1367/7-1 (525826480), CR 811/3-1 (525826480) and supported by its central service project (ME 2210/11-1). HG is grateful for the computational infrastructure and support provided by the “Zentrum für Informations-und Medientechnologie” (ZIM) at Heinrich Heine University Düsseldorf and the computing time provided by the John von Neumann Institute for Computing (NIC) on the supercomputers JUWELS and JUPITER at Jülich Supercomputing Centre (JSC) (user ID: chitin). We thank the Cytometry and Cell Sorting Core Facility of the University Medical Center Hamburg-Eppendorf (UKE) for support.

## Ethics statement

The authors have nothing to report.

## Data availability statement

The data that support the findings of this study are available from the corresponding author upon request.

## References

1 No, H. K. & Meyers, S. P. Application of chitosan for treatment of wastewaters. Rev Environ Contam Toxicol 163, 1–27 (2000). 10.1007/978-1-4757-6429-1_1

2 Delic, M., Butorova, I. & Kuskov, A. Chitosan application in cosmetics and dermatology - Antimicrobial and prebiotic potential to control human microbiome. J Biotechnol 408, 217–231 (2025). 10.1016/j.jbiotec.2025.09.014

3 Mohan, K. et al. Chitin, chitosan and chitooligosaccharides as potential growth promoters and immunostimulants in aquaculture: A comprehensive review. Int J Biol Macromol 251, 126285 (2023). 10.1016/j.ijbiomac.2023.126285

4 Shigemasa, Y. & Minami, S. Applications of chitin and chitosan for biomaterials. Biotechnol Genet Eng Rev 13, 383–420 (1996). 10.1080/02648725.1996.10647935

5 Fuchs, K. et al. The fungal ligand chitin directly binds TLR2 and triggers inflammation dependent on oligomer size. EMBO Rep 19 (2018). 10.15252/embr.201846065

6 He, X. et al. LYSMD3: A mammalian pattern recognition receptor for chitin. Cell Rep 36, 109392 (2021). 10.1016/j.celrep.2021.109392

7 Moeller, J. B. et al. Modulation of the fungal mycobiome is regulated by the chitin-binding receptor FIBCD1. J Exp Med 216, 2689–2700 (2019). 10.1084/jem.20182244

8 Kawai, T., Ikegawa, M., Ori, D. & Akira, S. Decoding Toll-like receptors: Recent insights and perspectives in innate immunity. Immunity 57, 649–673 (2024). 10.1016/j.immuni.2024.03.004

9 Gorzelanny, C., Poppelmann, B., Pappelbaum, K., Moerschbacher, B. M. & Schneider, S. W. Human macrophage activation triggered by chitotriosidase-mediated chitin and chitosan degradation. Biomaterials 31, 8556–8563 (2010). 10.1016/j.biomaterials.2010.07.100

10 Hellmann, M. J. et al. Chitosan production methods influence receptor-mediated immune responses but not target-mediated antimicrobial bioactivities. Carbohydr Polym 371, 124487 (2026). 10.1016/j.carbpol.2025.124487

11 Hellmann, M. J. et al. Heterogeneously deacetylated chitosans possess an unexpected regular pattern favoring acetylation at every third position. Nat Commun 15, 6695 (2024). 10.1038/s41467-024-50857-1

12 Di Rosa, M., Distefano, G., Zorena, K. & Malaguarnera, L. Chitinases and immunity: Ancestral molecules with new functions. Immunobiology 221, 399–411 (2016). 10.1016/j.imbio.2015.11.014

13 Kwak, E. J. et al. Chitinase 3-like 1 drives allergic skin inflammation via Th2 immunity and M2 macrophage activation. Clin Exp Allergy 49, 1464–1474 (2019). 10.1111/cea.13478

14 Harvey, S., Whaley, J. & Eberhardt, K. The relationship between serum levels of YKL-40 and disease progression in patients with early rheumatoid arthritis. Scand J Rheumatol 29, 391–393 (2000). 10.1080/030097400447606

15 Koutroubakis, I. E. et al. Increased serum levels of YKL-40 in patients with inflammatory bowel disease. Int J Colorectal Dis 18, 254–259 (2003). 10.1007/s00384-002-0446-z

16 Geng, B. et al. Chitinase 3-like 1-CD44 interaction promotes metastasis and epithelial-to-mesenchymal transition through beta-catenin/Erk/Akt signaling in gastric cancer. J Exp Clin Cancer Res 37, 208 (2018). 10.1186/s13046-018-0876-2

17 von Palubitzki, L. et al. Differences of the tumour cell glycocalyx affect binding of capsaicin-loaded chitosan nanocapsules. Sci Rep 10, 22443 (2020). 10.1038/s41598-020-79882-y

18 Clausen, T. M. et al. Antithrombin-binding heparan sulfate is ubiquitously expressed in epithelial cells and suppresses pancreatic tumorigenesis. J Clin Invest 135 (2025). 10.1172/JCI184172

19 Gyimesi, M., Okolicsanyi, R. K. & Haupt, L. M. Beyond amyloid and tau: rethinking Alzheimer’s disease through less explored avenues. Open Biol 14, 240035 (2024). 10.1098/rsob.240035

20 Patterson, A. M. et al. Expression of heparan sulfate proteoglycans in murine models of experimental colitis. Inflamm Bowel Dis 18, 1112–1126 (2012). 10.1002/ibd.21879

21 Annaval, T. et al. Heparan Sulfate Proteoglycans Biosynthesis and Post Synthesis Mechanisms Combine Few Enzymes and Few Core Proteins to Generate Extensive Structural and Functional Diversity. Molecules 25 (2020). 10.3390/molecules25184215

22 Vives, R. R., Seffouh, A. & Lortat-Jacob, H. Post-Synthetic Regulation of HS Structure: The Yin and Yang of the Sulfs in Cancer. Front Oncol 3, 331 (2014). 10.3389/fonc.2013.00331

23 Hellmann, M. J., Moerschbacher, B. M. & Cord-Landwehr, S. Fast insights into chitosan-cleaving enzymes by simultaneous analysis of polymers and oligomers through size exclusion chromatography. Scientific Reports 14, 3417 (2024). 10.1038/s41598-024-54002-2

24 Kosters, M. et al. pymzML v2.0: introducing a highly compressed and seekable gzip format. Bioinformatics 34, 2513–2514 (2018). 10.1093/bioinformatics/bty046

25 Cord-Landwehr, S. et al. Quantitative Mass-Spectrometric Sequencing of Chitosan Oligomers Revealing Cleavage Sites of Chitosan Hydrolases. Anal Chem 89, 2893–2900 (2017). 10.1021/acs.analchem.6b04183

26 Houston, D. R., Recklies, A. D., Krupa, J. C. & van Aalten, D. M. Structure and ligand-induced conformational change of the 39-kDa glycoprotein from human articular chondrocytes. J Biol Chem 278, 30206–30212 (2003). 10.1074/jbc.M303371200

27 Sastry, G. M., Adzhigirey, M., Day, T., Annabhimoju, R. & Sherman, W. Protein and ligand preparation: parameters, protocols, and influence on virtual screening enrichments. J Comput Aided Mol Des 27, 221–234 (2013). 10.1007/s10822-013-9644-8

28 Delas, T. et al. Effects of Chain Length of Chitosan Oligosaccharides on Solution Properties and Complexation with siRNA. Polymers (Basel) 11 (2019). 10.3390/polym11081236

29 Roe, D. R. & Bergonzo, C. prepareforleap: An automated tool for fast PDB-to-parameter generation. J Comput Chem 43, 930–935 (2022). 10.1002/jcc.26847

30 Izadi, S., Anandakrishnan, R. & Onufriev, A. V. Building Water Models: A Different Approach. J Phys Chem Lett 5, 3863–3871 (2014). 10.1021/jz501780a

31 Machado, M. R. & Pantano, S. Split the Charge Difference in Two! A Rule of Thumb for Adding Proper Amounts of Ions in MD Simulations. J Chem Theory Comput 16, 1367–1372 (2020). 10.1021/acs.jctc.9b00953

32 D.A. Case, H. M. A., K. Belfon, I.Y. Ben-Shalom, J.T. Berryman, S.R. Brozell, D.S. Cerutti, T.E., Cheatham, I., G.A. Cisneros, V.W.D. Cruzeiro, T.A. Darden, N. Forouzesh, M. Ghazimirsaeed, G. Gi-, ambasu, T. G., M.K. Gilson, H. Gohlke, A.W. Goetz, J. Harris, Z. Huang, S. Izadi, S.A. Izmailov, K., Kasavajhala, M. C. K., I. Kolossváry, A. Kovalenko, T. Kurtzman, T.S. Lee, P. Li, Z. Li, C. Lin, J. Liu,, T. Luchko, R. L., M. Machado, M. Manathunga, K.M. Merz, Y. Miao, O. Mikhailovskii, G. Monard, H., Nguyen, K. A. O. H., A. Onufriev, F. Pan, S. Pantano, A. Rahnamoun, D.R. Roe, A. Roitberg, C. Sagui,, S. Schott-Verdugo, A. S., J. Shen, C.L. Simmerling, N.R. Skrynnikov, J. Smith, J. Swails, R.C. Walker,, & J. Wang, J. W., X. Wu, Y. Wu, Y. Xiong, Y. Xue, D.M. York, C. Zhao, Q. Zhu, and P.A. Kollman. Amber 2024. (University of California, San Francisco, 2024).

33 Tian, C. et al. ff19SB: Amino-Acid-Specific Protein Backbone Parameters Trained against Quantum Mechanics Energy Surfaces in Solution. J Chem Theory Comput 16, 528–552 (2020). 10.1021/acs.jctc.9b00591

34 Kirschner, K. N. et al. GLYCAM06: a generalizable biomolecular force field. Carbohydrates. J Comput Chem 29, 622–655 (2008). 10.1002/jcc.20820

35 Li, P., Song, L. F. & Merz, K. M., Jr. Systematic Parameterization of Monovalent Ions Employing the Nonbonded Model. J Chem Theory Comput 11, 1645–1657 (2015). 10.1021/ct500918t

36 Sengupta, A., Li, Z., Song, L. F., Li, P. & Merz, K. M., Jr. Parameterization of Monovalent Ions for the OPC3, OPC, TIP3P-FB, and TIP4P-FB Water Models. *J Chem Inf Model* 61, 869-880 (2021). 10.1021/acs.jcim.0c01390

37 Le Grand, S., Götz, A. W. & Walker, R. C. SPFP: Speed without compromise—A mixed precision model for GPU accelerated molecular dynamics simulations. Computer Physics Communications 184, 374–380 (2013). 10.1016/j.cpc.2012.09.022

38 Salomon-Ferrer, R., Gotz, A. W., Poole, D., Le Grand, S. & Walker, R. C. Routine Microsecond Molecular Dynamics Simulations with AMBER on GPUs. 2. Explicit Solvent Particle Mesh Ewald. J Chem Theory Comput 9, 3878–3888 (2013). 10.1021/ct400314y

39 Yuksel, S. et al. Uncoupling of Voltage-and Ligand-Induced Activation in HCN2 Channels by Glycine Inserts. Front Physiol 13, 895324 (2022). 10.3389/fphys.2022.895324

40 Berendsen, H. J. C., Postma, J. P. M., van Gunsteren, W. F., DiNola, A. & Haak, J. R. Molecular dynamics with coupling to an external bath. The Journal of Chemical Physics 81, 3684–3690 (1984). 10.1063/1.448118

41 Loncharich, R. J., Brooks, B. R. & Pastor, R. W. Langevin dynamics of peptides: the frictional dependence of isomerization rates of N-acetylalanyl-N’-methylamide. Biopolymers 32, 523–535 (1992). 10.1002/bip.360320508

42 Ryckaert, J.-P., Ciccotti, G. & Berendsen, H. J. C. Numerical integration of the cartesian equations of motion of a system with constraints: molecular dynamics of n-alkanes. Journal of Computational Physics 23, 327–341 (1977). 10.1016/0021-9991(77)90098-5

43 Darden, T., York, D. & Pedersen, L. Particle mesh Ewald: An N⋅log(N) method for Ewald sums in large systems. The Journal of Chemical Physics 98, 10089–10092 (1993). 10.1063/1.464397

44 Hopkins, C. W., Le Grand, S., Walker, R. C. & Roitberg, A. E. Long-Time-Step Molecular Dynamics through Hydrogen Mass Repartitioning. J Chem Theory Comput 11, 1864–1874 (2015). 10.1021/ct5010406

45 Roe, D. R. & Cheatham, T. E., 3rd. PTRAJ and CPPTRAJ: Software for Processing and Analysis of Molecular Dynamics Trajectory Data. J Chem Theory Comput 9, 3084–3095 (2013). 10.1021/ct400341p

46 Humphrey, W., Dalke, A. & Schulten, K. VMD: Visual molecular dynamics. Journal of Molecular Graphics 14, 33–38 (1996). 10.1016/0263-7855(96)00018-5

47 Gohlke, H. & Case, D. A. Converging free energy estimates: MM-PB(GB)SA studies on the protein–protein complex Ras–Raf. Journal of Computational Chemistry 25, 238–250 (2004). 10.1002/jcc.10379

48 Homeyer, N. & Gohlke, H. Free Energy Calculations by the Molecular Mechanics Poisson−Boltzmann Surface Area Method. Molecular Informatics 31, 114–122 (2012). 10.1002/minf.201100135

49 Miller, B. R., III et al. MMPBSA.py: An Efficient Program for End-State Free Energy Calculations. Journal of Chemical Theory and Computation 8, 3314–3321 (2012). 10.1021/ct300418h

50 Homeyer, N., Stoll, F., Hillisch, A. & Gohlke, H. Binding Free Energy Calculations for Lead Optimization: Assessment of Their Accuracy in an Industrial Drug Design Context. Journal of Chemical Theory and Computation 10, 3331–3344 (2014). 10.1021/ct5000296

51 Hou, T., Wang, J., Li, Y. & Wang, W. Assessing the Performance of the MM/PBSA and MM/GBSA Methods. 1. The Accuracy of Binding Free Energy Calculations Based on Molecular Dynamics Simulations. Journal of Chemical Information and Modeling 51, 69–82 (2011). 10.1021/ci100275a

52 Masukawa, K. M., Kollman, P. A. & Kuntz, I. D. Investigation of Neuraminidase-Substrate Recognition Using Molecular Dynamics and Free Energy Calculations. Journal of Medicinal Chemistry 46, 5628–5637 (2003). 10.1021/jm030060q

53 Sun, H. et al. Assessing the performance of MM/PBSA and MM/GBSA methods. 7. Entropy effects on the performance of end-point binding free energy calculation approaches. Physical Chemistry Chemical Physics 20, 14450–14460 (2018). 10.1039/C7CP07623A

54 Virtanen, P. et al. SciPy 1.0: fundamental algorithms for scientific computing in Python. Nature Methods 17, 261–272 (2020). 10.1038/s41592-019-0686-2

55 Tyrikos-Ergas, T. et al. Systematic Structural Characterization of Chitooligosaccharides Enabled by Automated Glycan Assembly. Chemistry – A European Journal 27, 2321–2325 (2021). 10.1002/chem.202005228

56 Nampally, M., Moerschbacher, B. M. & Kolkenbrock, S. Fusion of a novel genetically engineered chitosan affinity protein and green fluorescent protein for specific detection of chitosan in vitro and in situ. Appl Environ Microbiol 78, 3114–3119 (2012). 10.1128/AEM.07506-11

57 Grossdorf, A., Obser, T., Wang, Y. & Gorzelanny, C. Single Molecule Force Spectroscopy at Cell Surfaces to Study Physical Properties of Heparan Sulfate Chains and Protein-Heparan Sulfate Interactions. Proteoglycan Research 3, e70017 (2025). 10.1002/pgr2.70017

58 Wang, Y. et al. Heparan sulfate dependent binding of plasmatic von Willebrand factor to blood circulating melanoma cells attenuates metastasis. Matrix Biol 111, 76–94 (2022). 10.1016/j.matbio.2022.06.002

59 Evans, E. & Ritchie, K. Dynamic strength of molecular adhesion bonds. Biophys J 72, 1541–1555 (1997). 10.1016/S0006-3495(97)78802-7

60 Gude, F. et al. Hedgehog is relayed through dynamic heparan sulfate interactions to shape its gradient. Nat Commun 14, 758 (2023). 10.1038/s41467-023-36450-y

61 Sembajwe, L. F., Katta, K., Gronning, M. & Kusche-Gullberg, M. The exostosin family of glycosyltransferases: mRNA expression profiles and heparan sulphate structure in human breast carcinoma cell lines. Biosci Rep 38 (2018). 10.1042/BSR20180770

62 Richter, C., Cord-Landwehr, S., Singh, R., Ryll, J. & Moerschbacher, B. M. Dissecting and optimizing bioactivities of chitosans by enzymatic modification. Carbohydrate Polymers 349, 122958 (2025). 10.1016/j.carbpol.2024.122958

63 Fusetti, F. et al. Structure of human chitotriosidase. Implications for specific inhibitor design and function of mammalian chitinase-like lectins. J Biol Chem 277, 25537–25544 (2002). 10.1074/jbc.M201636200

64 Hellmann, M. J., Marongiu, G. L., Gorzelanny, C., Moerschbacher, B. M. & Cord-Landwehr, S. Hydrolysis of chitin and chitosans by the human chitinolytic enzymes: chitotriosidase, acidic mammalian chitinase, and lysozyme. Int J Biol Macromol 297, 139789 (2025). 10.1016/j.ijbiomac.2025.139789

65 Metz, A. et al. Hot Spots and Transient Pockets: Predicting the Determinants of Small-Molecule Binding to a Protein–Protein Interface. Journal of Chemical Information and Modeling 52, 120–133 (2012). 10.1021/ci200322s

66 Kognole, A. A. & Payne, C. M. Inhibition of Mammalian Glycoprotein YKL-40: IDENTIFICATION OF THE PHYSIOLOGICAL LIGAND. J Biol Chem 292, 2624–2636 (2017). 10.1074/jbc.M116.764985

67 Ngernyuang, N. et al. A Heparin Binding Motif Rich in Arginine and Lysine is the Functional Domain of YKL-40. Neoplasia 20, 182–192 (2018). 10.1016/j.neo.2017.11.011

68 Mosier, P. D., Krishnasamy, C., Kellogg, G. E. & Desai, U. R. On the specificity of heparin/heparan sulfate binding to proteins. Anion-binding sites on antithrombin and thrombin are fundamentally different. PLoS One 7, e48632 (2012). 10.1371/journal.pone.0048632

69 Zhang, F. et al. Potential Anti-SARS-CoV-2 Activity of Pentosan Polysulfate and Mucopolysaccharide Polysulfate. Pharmaceuticals (Basel) 15 (2022). 10.3390/ph15020258

70 van den Born, J., et al. Novel heparan sulfate structures revealed by monoclonal antibodies. J Biol Chem 280, 20516–20523 (2005). 10.1074/jbc.M502065200

71 Koch, A., Bonus, M., Gohlke, H. & Klöcker, N. Isoform-specific Inhibition of N-methyl-D-aspartate Receptors by Bile Salts. Scientific Reports 9, 10068 (2019). 10.1038/s41598-019-46496-y

72 Sorlier, P., Rochas, C., Morfin, I., Viton, C. & Domard, A. Light scattering studies of the solution properties of chitosans of varying degrees of acetylation. Biomacromolecules 4, 1034–1040 (2003). 10.1021/bm034054n

73 Sorlier, P., Denuziere, A., Viton, C. & Domard, A. Relation between the degree of acetylation and the electrostatic properties of chitin and chitosan. Biomacromolecules 2, 765–772 (2001). 10.1021/bm015531+

74 Schatz, C., Viton, C., Delair, T., Pichot, C. & Domard, A. Typical physicochemical behaviors of chitosan in aqueous solution. Biomacromolecules 4, 641–648 (2003). 10.1021/bm025724c

75 Shao, R. et al. YKL-40, a secreted glycoprotein, promotes tumor angiogenesis. Oncogene 28, 4456–4468 (2009). 10.1038/onc.2009.292

76 Szydlowski, M. et al. PIM kinase inhibition attenuates pro-tumoral and immunosuppressive functions of macrophages in classic Hodgkin lymphoma. Cell Death Dis 17, 136 (2025). 10.1038/s41419-025-08402-5

77 Kim, S. et al. Carcinoma-produced factors activate myeloid cells through TLR2 to stimulate metastasis. Nature 457, 102–106 (2009). 10.1038/nature07623

78 Martinez-Ordonez, A. et al. Hyaluronan driven by epithelial aPKC deficiency remodels the microenvironment and creates a vulnerability in mesenchymal colorectal cancer. Cancer Cell 41, 252–271 e259 (2023). 10.1016/j.ccell.2022.11.016

79 Dominguez-Gutierrez, P. R. et al. Hyal2 Expression in Tumor-Associated Myeloid Cells Mediates Cancer-Related Inflammation in Bladder Cancer. Cancer Res 81, 648–657 (2021). 10.1158/0008-5472.CAN-20-1144

80 Witschen, P. M. et al. Tumor Cell Associated Hyaluronan-CD44 Signaling Promotes Pro-Tumor Inflammation in Breast Cancer. Cancers (Basel) 12 (2020). 10.3390/cancers12051325

81 Sokolowska, M. et al. Low molecular weight hyaluronan activates cytosolic phospholipase A2alpha and eicosanoid production in monocytes and macrophages. J Biol Chem 289, 4470–4488 (2014). 10.1074/jbc.M113.515106

82 Swart, M. et al. The extracellular heparan sulfatase SULF2 limits myeloid IFNbeta signaling and Th17 responses in inflammatory arthritis. Cell Mol Life Sci 81, 350 (2024). 10.1007/s00018-024-05333-w

83 Johansen, J. S. et al. Serum YKL-40 concentrations in patients with rheumatoid arthritis: relation to disease activity. Rheumatology (Oxford) 38, 618–626 (1999). 10.1093/rheumatology/38.7.618

84 Deng, Y. et al. Serum CHI3L1 levels correlate with disease activity in rheumatoid arthritis and reveal potential molecular mechanisms. Front Immunol 16, 1729989 (2025). 10.3389/fimmu.2025.1729989

85 Magnusdottir, U. et al. Heparin-binding of the human chitinase-like protein YKL-40 is allosterically modified by chitin oligosaccharides. Biochem Biophys Rep 41, 101908 (2025). 10.1016/j.bbrep.2024.101908

86 Wright, S. D., Ramos, R. A., Tobias, P. S., Ulevitch, R. J. & Mathison, J. C. CD14, a receptor for complexes of lipopolysaccharide (LPS) and LPS binding protein. Science 249, 1431–1433 (1990). 10.1126/science.1698311

87 Poltorak, A. et al. Defective LPS signaling in C3H/HeJ and C57BL/10ScCr mice: mutations in Tlr4 gene. Science 282, 2085–2088 (1998). 10.1126/science.282.5396.2085

88 Wurfel, M. M. et al. Targeted deletion of the lipopolysaccharide (LPS)-binding protein gene leads to profound suppression of LPS responses ex vivo, whereas in vivo responses remain intact. J Exp Med 186, 2051–2056 (1997). 10.1084/jem.186.12.2051

89 Zheng, Q. et al. Decoding the CHI3L1/IL-13Ralpha2 signaling nexus in MASH-fibrosis pathogenesis. Sci Adv 11, eadz3223 (2025). 10.1126/sciadv.adz3223

