## Supplemental Figures for "Chitinase-3-like protein 1 decodes chitosan acetylation patterns into toll-like receptor 2 signaling through heparan sulfate"

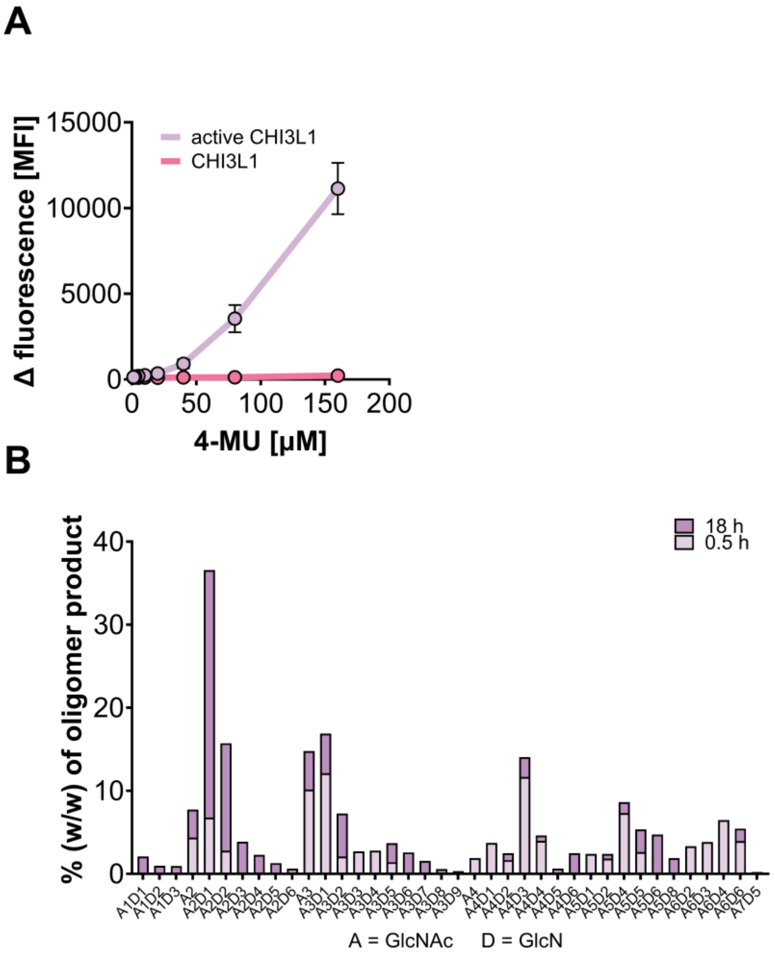

**Supplementary Fig. 1: Enzymatic activity and mass spectrometry analysis of chitosan digestion products generated by active CHI3L1 (A138D/L140E).**

**A:** Cleavage of 4-methylumbelliferone chitobioside substrate (4-MU) by active CHI3L1. Wild-type CHI3L1 served as control. 4-MU substrate was cleaved by active CHI3L1, whereas no cleavage was observed for CHI3L1.

**B:** Chitin and chitosan oligosaccharides generated after incubation of chitosan (DA 48%) with activated CHI3L1 for 0.5 h and 18 h. Oligomer composition is shown as percentage (w/w) of total oligomer products. A = GlcNAc; D = GlcN.

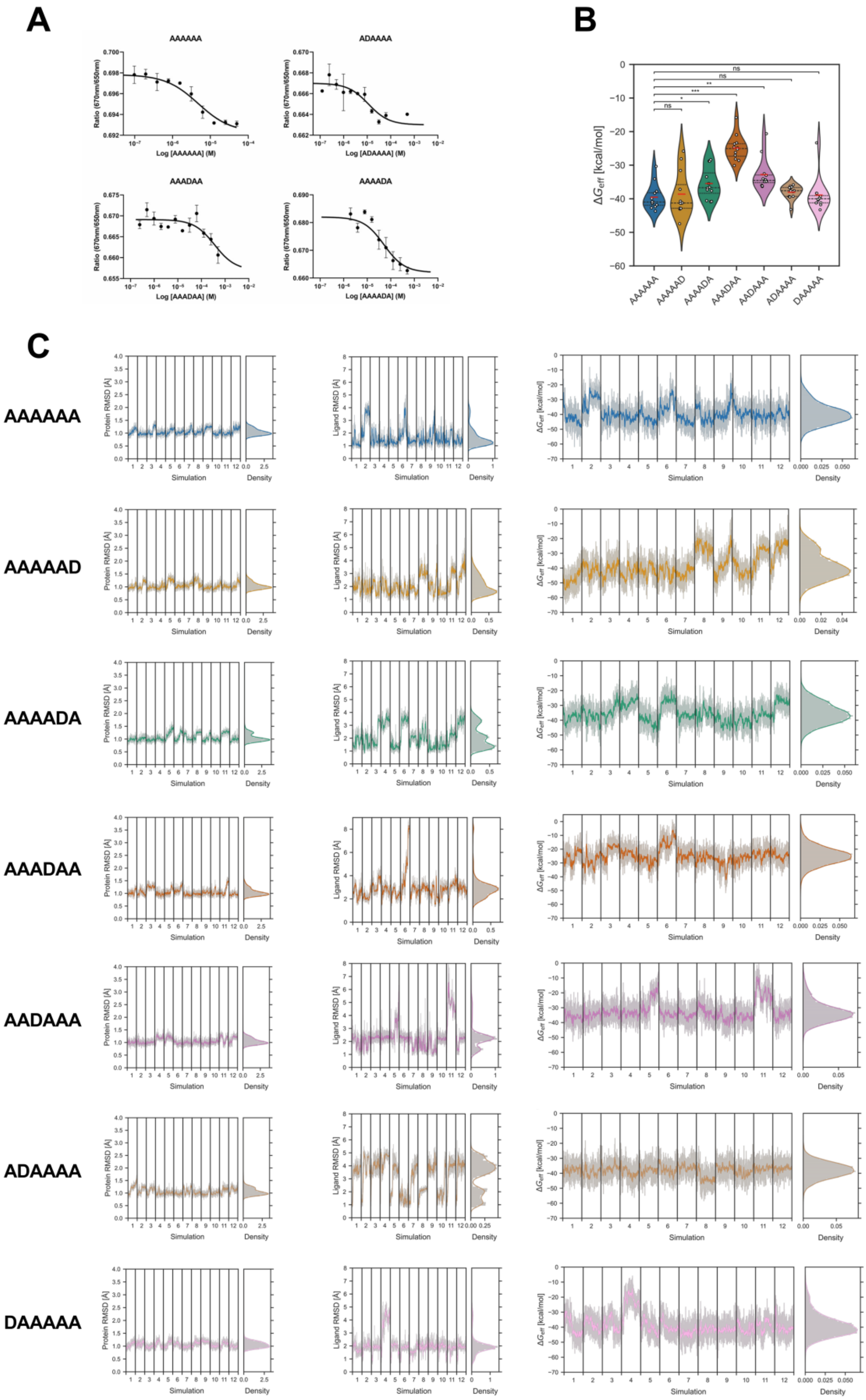

**Supplementary Fig. 2: Complementary microscale thermophoresis and molecular dynamics simulation analysis of chitin hexamers with different PA binding to CHI3L1.**

**A:** MST analysis of chitin oligosaccharide binding to CHI3L1. Binding isotherms obtained from the titration of chemically-defined COS to fluorescently labeled CHI3L1. Error estimates are given as 68% confidence intervals from three independent experiments.

**B:** Effective binding energy of COS binding to CHI3L1. The violin plot shows the distribution of the effective binding energies of the COS derivatives to CHI3L1. The interquartile range is bounded by the dotted lines, while the median is represented by the dashed line. The red line represents the mean. The statistical significance between the mean values of the partially acetylated COS and the fully acetylated ligand (AAAAAA) was evaluated using a two-sided Welch's t-test (ns: not significant; *: $p < 0.05$, **: $p < 0.01$, ***: $p < 0.001$).

**C:** Structural and energetic variability in the CHI3L1- COS complexes during the MD simulations. Time courses and probability distributions of the RMSD of CHI3L1 (Cα atoms, **left panels**), COS ligands (heavy atoms, **middle panels**), and effective binding energies ($\Delta G_{eff}$, **right panels**), calculated using the MM-PBSA method with an internal dielectric constant of $\varepsilon int = 4$, shown for all of the 12 simulations (1 µs length). Thick, opaque lines represent the data smoothed with a Savitzky-Golay filter (window length = 51, degree of the smoothing polynomial = 2); thin, transparent lines show the unsmoothed data. The smoothed distributions (thick, opaque lines) were calculated using a Gaussian kernel density estimator (bandwidth determined via Scott's rule). The underlying data is displayed as a histogram (transparent bars). RMSD values were calculated with respect to the crystal structure (PDB ID: 1HJW).

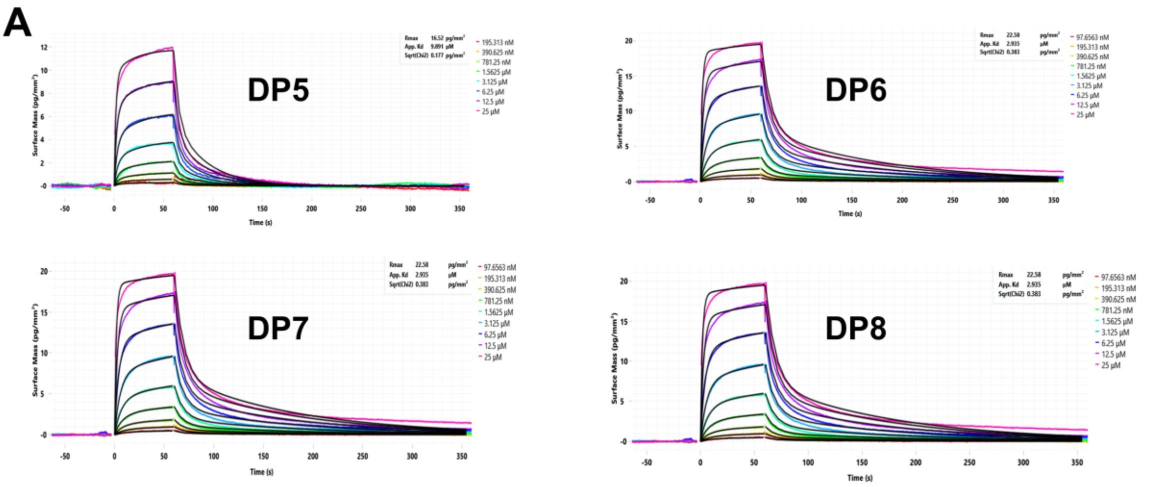

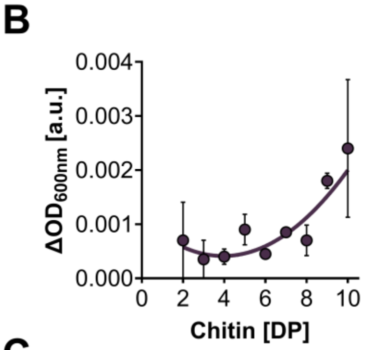

C

| name | production | average DP | SD DP | average Mw [kDa] | SD Mw | average DA | SD DA | authors | doi |
|---|---|---|---|---|---|---|---|---|---|
| C50 | CNA | 2518 | 85 | 457 | 1 | 49.5 | | G. Lamarque, J.M. Lucas, C. Viton, A. Domard | 10.1021/bm0496357 |
| C50 | CNA | 1024 | 42 | 182.4 | 7.48 | 40.9 | 0.22 | K. Eickelpasch, L. Brussel, S. Cord-Landwehr, B. M. Moerschbacher, C. Richter | 10.1016/j.carbpol.2026.125404 |
| C40 | CNA | 1178 | | 209.4 | 0 | 040 | | | |
| C32 | CNA | 1027 | 6 | 177 | 1.03 | 27.04 | 0.15 | K. Eickelpasch, L. Brussel, S. Cord-Landwehr, B. M. Moerschbacher, C. Richter | 10.1016/j.carbpol.2026.125404 |
| C24 | CNA | 1389 | 81 | 238 | 14 | 24.4 | 0.3 | M. J. Hellmann, K. Eickelpasch, A. Großdorf, P. Barreto, M. Schwarzländer, C. Gorzelanny, B. M. Moerschbacher, C. Richter | 10.1016/j.carbpol.2025.124487 |
| H24 | HTDA | 677 | 0 | 116 | 0 | 24.2 | 0.4 | M. J. Hellmann, K. Eickelpasch, A. Großdorf, P. Barreto, M. Schwarzländer, C. Gorzelanny, B. M. Moerschbacher, C. Richter | 10.1016/j.carbpol.2025.124487 |
| C15 | CNA | 1113 | 4 | 185.4 | 0.66 | 13.35 | 0.4 | K. Eickelpasch, L. Brussel, S. Cord-Landwehr, B. M. Moerschbacher, C. Richter | 10.1016/j.carbpol.2026.125404 |
| C10 | CNA | 2580 | 79 | 426 | 1 | 9.8 | | G. Lamarque, J.M. Lucas, C. Viton, A. Domard | 10.1021/bm0496357 |
| C0 | CNA | 1237 | 21 | 199.4 | 3.39 | 0.56 | 0.03 | K. Eickelpasch, L. Brussel, S. Cord-Landwehr, B. M. Moerschbacher, C. Richter | 10.1016/j.carbpol.2026.125404 |

**Supplementary Fig. 3: Characterization of CHI3L1 and mutants in chitin binding.**
**A:** Representative GCI sensorgrams showing the binding of chitin oligomers (DP 5-8) to surface immobilized CHI3L1 at different concentrations. Surface mass is plotted as a function of time.
**B:** Light scattering of chitin oligomers (DP2-10) measured at 600 nm. Increased scattering above DP8 indicates the formation of chitin aggregates. A second-degree polynomial fit was used to guide the eye.
**C:** Summary of all used chitosans and their physiochemical properties, including production methods, average degree of polymerization (DP), average molecular weight (Mw), degree of acetylation (DA) and references.

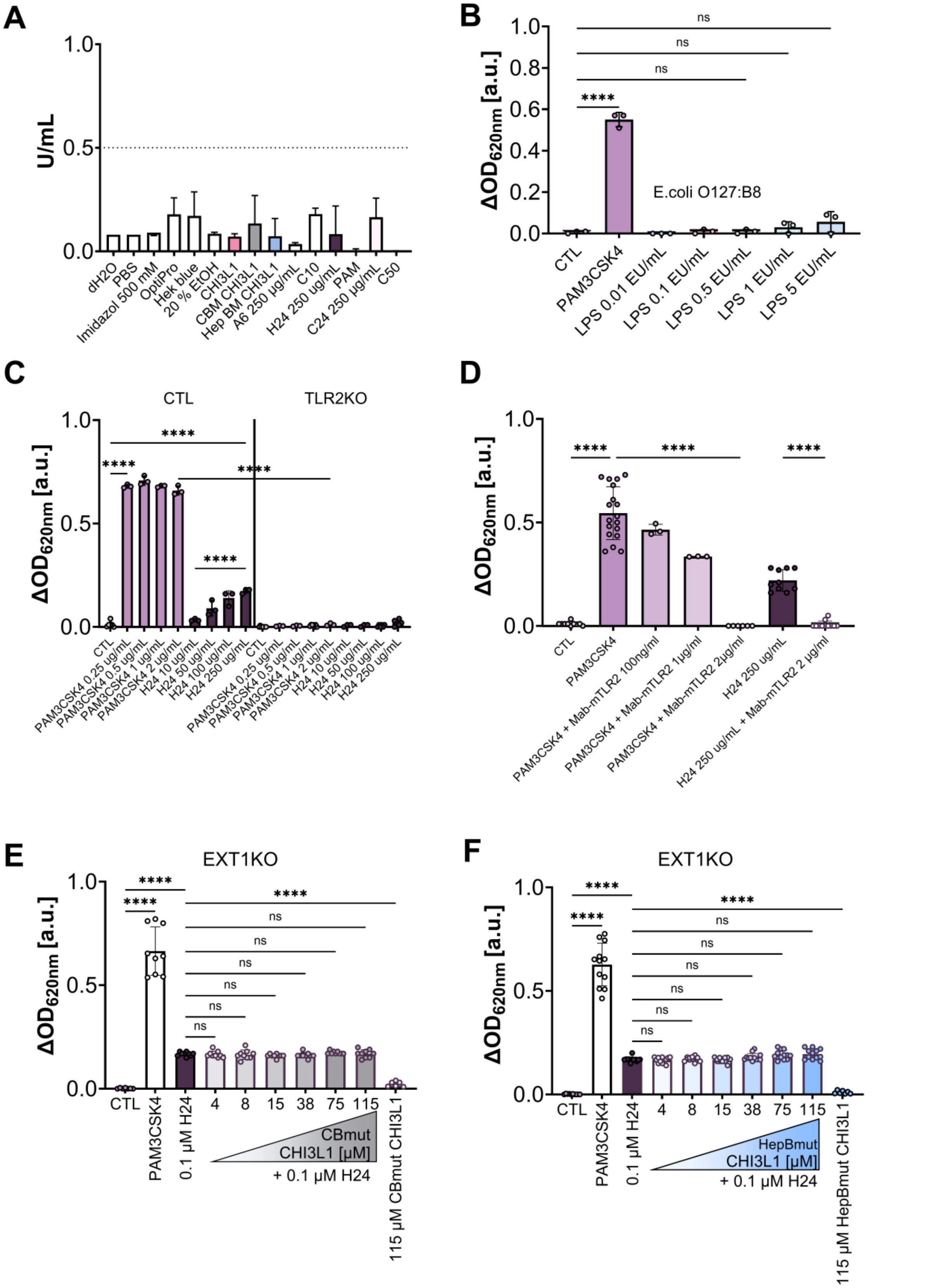

**Supplementary Fig. 4: Endotoxin controls and TLR2 and HS dependent cell activation.**

**A:** Endotoxin levels of all samples and solutions used for stimulation experiments. The dashed line indicates the critical endotoxins threshold of 0.5 EU/mL. All chitosans and solutions used were below this threshold.

**B:** HEK-Blue™ hTLR2 cells response to LPS (*E.coli* O127:B8) at the indicated concentrations. The cells showed slight response upon treatment with 5 EU/mL.
**C:** TLR2KO in HEK-Blue™ hTLR2 abolishes cellular response to PAM3CSK4 and H24. Control HEK-Blue™ hTLR2 cells expressing non-targeting gRNA were used as control.
**D:** Neutralization of TLR2 signalling through a TLR2 directed antibody (Mab-mTLR2) reduced the response to PAM3CSK4 and H24 in a dose dependent manner.
**E:** EXT1KO HEK-Blue™ hTLR2 cells stimulated with H24 and increasing concentration of CBmut CHI3L1. The CBmut CHI3L1 mutant failed to enhance signaling.
**F:** EXT1KO HEK-Blue™ hTLR2 cells stimulated with H24 and increasing concentration of HepBmut CHI3L1. Loss of EXT1 inhibits the CHI3L1-dependent enhancement of signaling, indicating that HS is required for TLR2-mediated cell activation.

Data represent mean ± SD, n = 3-5. Statistical analysis was performed using one-way ANOVA with Tukey' multiple-comparison test, * $P<0.05$; ** $P<0.01$; *** $P<0.001$; **** $P<0.0001$.

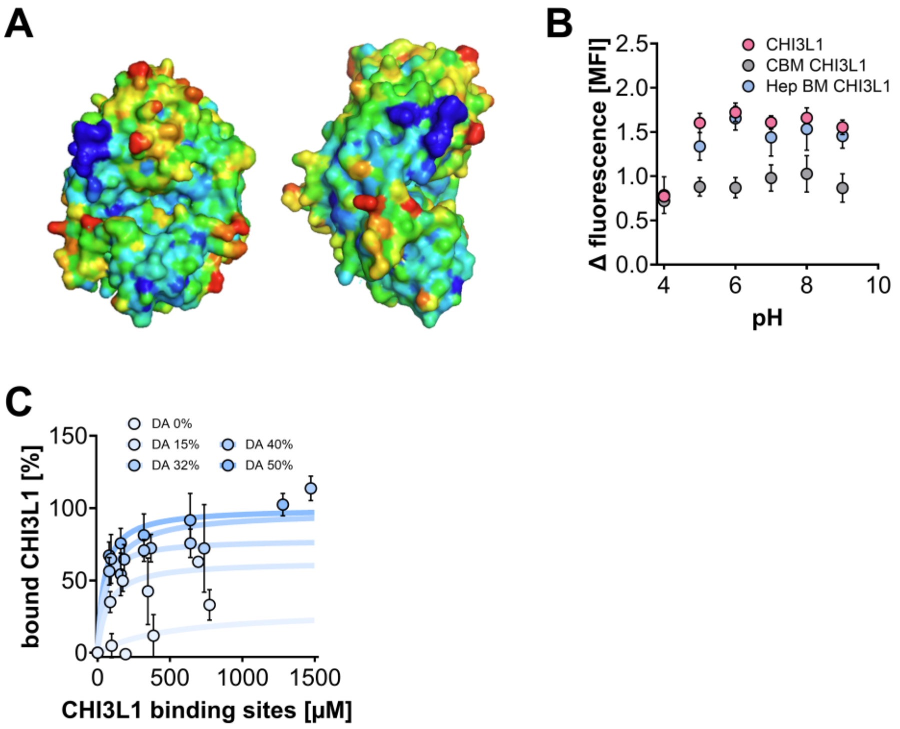

**D**

| block | name | length | structure | NS | 6S | 2S | 3S | total |
|---|---|---|---|---|---|---|---|---|
| 47 | G-76 | 12 | GlcNS6S-GlcA-GlcNS6S-GlcA- GlcNS6S-GlcA-GlcNS6S-GlcA-GlcNS6S-GlcA-GlcNS6S-GlcA-Az | 6 | 6 | 0 | 0 | 12 |
| 48 | 18Mer-1 | 18 | GlcNS6S-GlcA-GlcNS6S-GlcA- GlcNS6S-GlcA-GlcNS6S-GlcA-GlcNS6S-GlcA- GlcNS6S-GlcA-GlcNS6S-GlcA-GlcNS6S-GlcA-GlcNS6S-GlcA-Az | 9 | 9 | 0 | 0 | 18 |
| 69 | 75 | 9 | GlcA-GlcNS6S-GlcA-GlcNS6S-GlcA-GlcNS6S-IdoA2S-GlcNS6S-GlcA-Az | 4 | 4 | 1 | 0 | 9 |
| 74 | G-93 | 12 | GlcNS6S-GlcA-GlcNS6S-IdoA2S-GlcNS6S-IdoA2S-GlcNS6S-IdoA2S-GlcNS6S-IdoA2S-GlcNS6S-GlcA-Az | 6 | 6 | 4 | 0 | 16 |
| 76 | 14mer-2 | 14 | GlcNS6S-GlcA-GlcNS6S-IdoA2S-GlcNS6S-IdoA2S-GlcNS6S-IdoA2S-GlcNS6S-IdoA2S-GlcNS6S-IdoA2S-GlcNS6S-GlcA-Az | 7 | 7 | 5 | 0 | 19 |
| 77 | 18mer-2 | 18 | GlcNS-GlcA-GlcNS-IdoA2S-GlcNS-IdoA2S-GlcNS-IdoA2S-GlcNS-IdoA2S-GlcNS-IdoA2S-GlcNS-IdoA2S-GlcNS-IdoA2S-GlcNS-GlcA-Az | 9 | 0 | 7 | 0 | 16 |
| 57 | G-9 | 7 | GlcA-GlcNS6S-IdoA2S-GlcNS6S-IdoA2S-GlcNS6S-GlcA-Az | 3 | 3 | 2 | 0 | 8 |
| 81 | B-78 | 7 | GlcA-GlcNAc6S-GlcA-GlcNS6S-IdoA2S-GlcNS6S3S-GlcA-Az | 2 | 3 | 1 | 1 | 7 |
| 82 | 30 | 7 | GlcA-GlcNS6S-GlcA-GlcNS6S3S-IdoA2S-GlcNS6S-GlcA-Az | 3 | 3 | 1 | 1 | 8 |
| 83 | 124 | 8 | GlcNAc6S-GlcA-GlcNS6S3S-IdoA2S-GlcNS6S-IdoA2S-GlcNS6S-GlcA-Az | 3 | 4 | 2 | 1 | 10 |
| 72 | Az#76-NH2 | 9 | GlcA-GlcNS6S-GlcA-GlcNS6S-IdoA2S-GlcNS6S-IdoA2S-GlcNS6S-GlcA-Az | 4 | 4 | 2 | 0 | 10 |
| 61 | 50 | 8 | GlcNS6S-GlcA-GlcNS6S-GlcA-GlcNS6S-IdoA2S-GlcNS6S-GlcA-Az | 4 | 4 | 1 | 0 | 9 |
| 54 | 29 | 7 | GlcA-GlcNS6S-GlcA-GlcNS6S-IdoA2S-GlcNS6S-GlcA-Az | 3 | 3 | 1 | 0 | 7 |

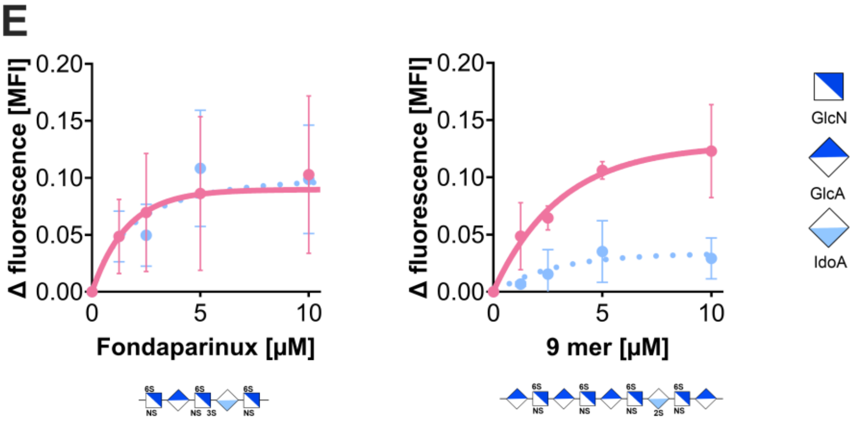

**Supplementary Fig. 5: HS binding site analysis and HepBmut CHI3L1.**

**A:** Schematic drawing of the B factor (Debye-Waller-Faktor) of CHI3L1 shown in two orientations. Red regions indicated higher structural flexibility, while blue regions indicated lower flexibility of the protein surface. The 1st HS binding site coincides with the most rigid region of CHI3L1.

**B** pH dependent chitin binding assay of CHI3L1 (pink), CBmut CHI3L1 (W31A/W99A/W352A, grey) and HepBmut CHI3l1 (R144A/R145A/K147A, blue) to chitin hexamer (A6). Proteins were excited with UV light (290 nm). Fluorescence emission was monitored at 340 nm. Binding remained stable between pH 5 and 9 for CHI3L1 and HepBmut CHI3L1 and decreased at pH 4. CBmut CHI3L1 showed no binding to A6.Data were normalized to buffer controls.

**C:** Change of the intrinsic Trp-fluorescence of HepBmut CHI3L1 by chitosans with increasing DA (0 – 50%), revealed stronger binding to high acetylated chitosans. CHI3L1 and HepBmut CHI3L1 exhibited same affinities to the selected chitosans. The chitosan binding of HepBmut CHI3L1 was not changed. Lines represent fits to a one-site specific binding model.

**D**: Summary of HS oligosaccharides testes in the glycan microarray that did not bind to HepB mut CHI3L1 compared to CHI3L1. Shown are the block number, name, chain length, monosaccharide sequence and sulfation pattern.

**E** Intrinsic Trp fluorescence analysis of CHI3L1 and HepBmut CHI3L1 binding to fondaparinux and 2S-6S 9-mer. Binding was weak and variable to fondaparinux but consistent to 2S-6S 9-mer. Binding of HepBmut CHI3L1 to 2S-6S 9-mer was reduced.

Data are shown as mean ± SD from n = 3-4 independent experiments. Statistical analysis was performed using a student's t test, *** P< 0.001; **** P <0.0001.

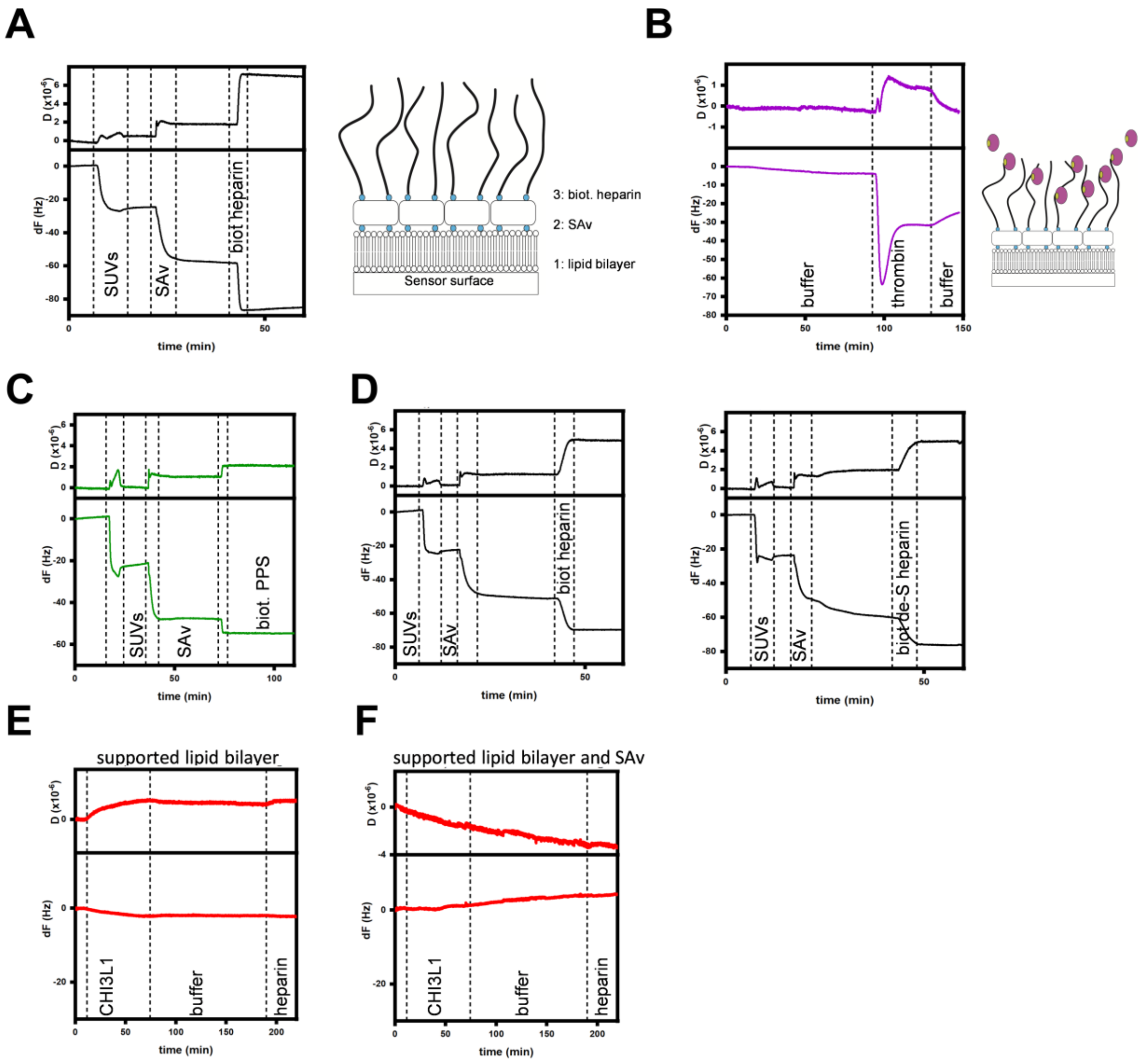

**Supplementary Fig. 6: QCM-D sensor assembly and control experiments.**

**A**: Functionalization of the sensor surface with heparin (**left**): SUVs: small unilamellar vesicles; SAv: streptavidin. Schematic overview of the functionalized chip surface (**right**): biotinylated lipid bilayer (SLB) with attached streptavidin (SAv) and biotinylated heparin to mimic cell surface HS.

**B**: Representative QCM-D sensogram of thrombin binding to HS functionalized surface. Showing characteristic increase in dissipation and parallel decrease in frequency associated with a single HS binding site. A schematic interaction of thrombin with HS functionalized surface is shown on the right side.

**C:** Functionalization of the sensor surface with pentosan polysulfate (PPS).

**D:** Functionalization of the sensor surface with heparin (**left**) and de-6S heparin (**right**).

**E:** Representative QCM-D sensogram of the interaction of CHI3L1 with the supported lipid bilayer on the chip surface as a negative control for possible unspecific interactions. No binding was observed.

**F:** Representative QCM-D sensogram of the interaction of CHI3L1 with the supported lipid bilayer and SAv on the chip surface as a negative control for possible unspecific interactions. No interaction was observed.

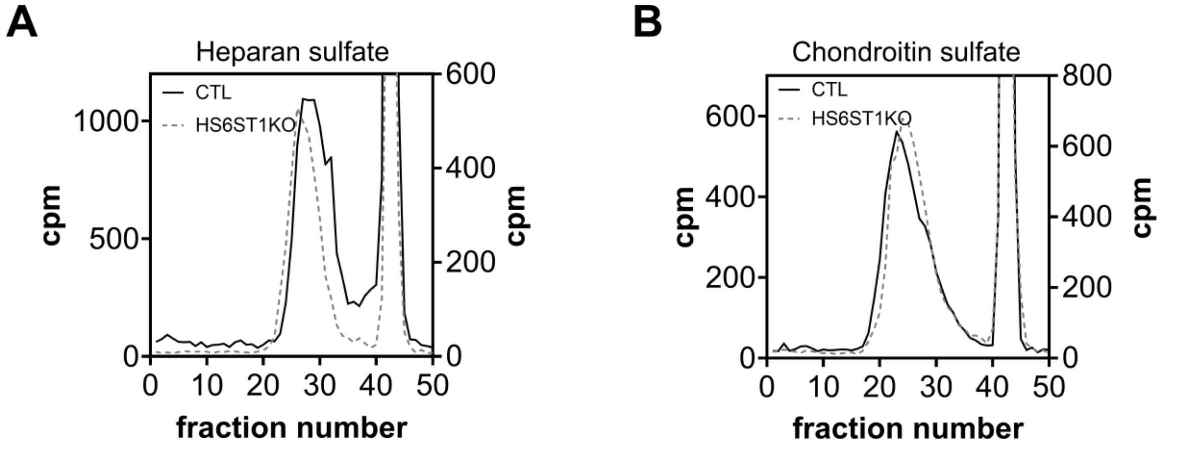

**Supplementary Fig. 7: Characterization of HS in HS6ST1KO B16F10 cells.**
**A:** Size-exclusion chromatography analysis of heparan sulfate isolated from control (CTL) and HS6ST1 KO cells. Lengths of the heparan chains of CTL and HS6ST1 cells were comparable. Shown are counts per minute (cpm) as a function of fraction number. The HS chain length was comparable between CTL and HS6ST1KO cells.
**B:** Size exclusion chromatography profiles of chondroitinsulfate isolated from CTL and HS6ST1KO B16F10 cells. The HS6ST1KO did not affect CS expression.

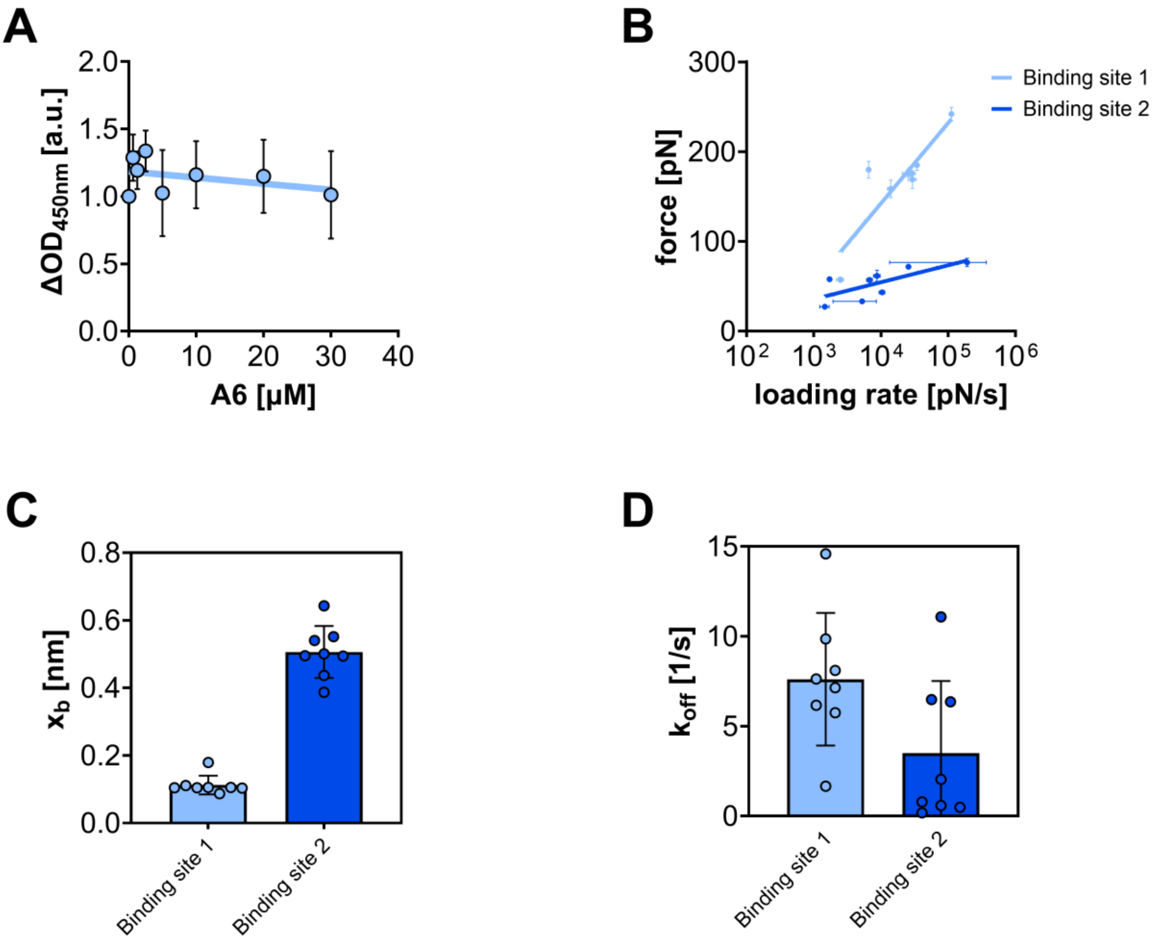

**Supplementary Fig. 8: Effect of chitin on CHI3L1-HS interaction.**

**A:** Heparin ELISA like binding assay demonstrating that chitin hexamer (A6) does not significantly affect the binding of HepBmut CHI3L1 to surface immobilized heparin at concentrations up to 30 µM. Lines represent fits to a one-site specific binding model.

**B:** SMFS experiments analysing the impact of chitin octamers on the interaction between heparin and CHI3L1. Shown are the measured rupture forces as a function of the applied loading rates in the presence of chitin octamer The linear fits are based on the Bell–Evans model and show different force dependencies for the two HS binding sites.

**C:** Depth of the energy barrier ($x_b$) and off rate ($k_{off}$) **(D)** for CHI3L1 obtained from the SMFS data (B). Compared to Figure 5E-J, presence of chitin had no impact on the interaction between HS binding and CHI3L1.
